# Multivalent mRNA tuberculosis vaccines protect against mycobacterial infection in zebrafish, mice and guinea pigs

**DOI:** 10.64898/2026.11.09.765257

**Authors:** Anni K Saralahti, Topias AA Isotalo, Mirja T Niskanen, Laura Kummola, Leena Mäkinen, Duc Nguyen, David F Ackart, Lauren Hunstad, Olivia R Miller, Ahmed Elkashif, Riley JK Easton, Martín González-Rodríguez, Maiju Junno, Susanna Valanne, Ilkka S Junttila, Brendan K Podell, Bernadette M Saunders, Mika Rämet

## Abstract

Despite a century of BCG use, tuberculosis remains a leading cause of death from infectious diseases and more effective vaccines are urgently needed. Here, we combined antigen discovery in zebrafish with mRNA vaccine technology to generate multivalent vaccine candidates, TUmRNA_4_, TUmRNA_6_ and TUmRNA_9_, encoding four, six, or nine *Mycobacterium tuberculosis* antigens, respectively. The vaccines were well tolerated and induced robust antigen-specific cellular immune responses, including IFNγ production and expansion of memory T-cell populations. In mice, TUmRNA_4_ and TUmRNA_6_ reduced pulmonary *M. tuberculosis* burden and lung pathology to levels comparable to BCG vaccination, and TUmRNA_6_ enhanced BCG-mediated protection when administered as a booster vaccine. In guinea pigs, TUmRNA_9_ reduced bacterial burden in the lungs, spleen, and lymph nodes and decreased pulmonary lesions. These findings demonstrate that protective antigens identified in zebrafish can be translated into effective multivalent mRNA vaccines and establish a framework for tuberculosis vaccine development across multiple preclinical models.

## Introduction

Tuberculosis remains a global health emergency and ongoing pandemic, especially affecting low- and middle-income countries^1^. In 2024, The World Health Organization (WHO) estimated that 10.7 million people developed tuberculosis and 1.23 million died from the disease, including 150,000 individuals living with HIV infection^1^. *Mycobacterium tuberculosis*, the primary causative agent of tuberculosis, is an airborne intracellular pathogen capable of surviving within macrophages by disrupting antimicrobial host processes, including phagosome maturation^2^. Although tuberculosis can be cured with prolonged antimicrobial treatment, the emergence of drug-resistant strains threatens progress in disease control^3–5^.

The Bacillus Calmette-Guérin (BCG) vaccine is a live attenuated strain of *Mycobacterium bovis* and is currently the only licensed and commercially available vaccine against tuberculosis. Approved for use in 1921, BCG is one of the most widely administered vaccines worldwide. After a century of clinical use it remains part of routine infant immunization programs in many countries, with over 100 million doses administered every year^6^. Although BCG protects young children against severe forms of tuberculosis, its protective efficacy begins to wane during adolescence^7^. In adults, who account for more than 90% of tuberculosis cases, BCG provides inconsistent protection against both primary infection and progression to active disease^8^. In addition, BCG vaccination can cause severe adverse effects, including BCG-associated osteitis/osteomyelitis and disseminated BCG infection, particularly in immunocompromised individuals^9–11^. Therefore, to stop the spread of tuberculosis, new vaccines that prevent primary infection, reduce the risk of disease reactivation, or enhance the protection conferred by BCG are needed.

Tuberculosis vaccine development has proven challenging, with no new EMA- or FDA-approved vaccines for clinical use since the introduction of BCG. This can largely be explained by the complex immunopathology of tuberculosis, limited commercial incentives in high-income countries, and the substantial financial investment required for lengthy clinical trials. Current vaccine candidates in late-stage development include the protein-subunit vaccine M72/AS01E^12^, live attenuated *M. tuberculosis* vaccine MTBVAC^13^, and the BCG-derived live attenuated VPM1002^14^, all of which are currently undergoing phase III clinical evaluation. In contrast, recent BCG revaccination studies have yielded limited efficacy, with no significant protection against sustained infection in HIV-negative adolescents tested negative for *M. tuberculosis*^15^.

Following the success of mRNA vaccines against COVID-19^16,17^, lipid nanoparticle (LNP) - encapsulated mRNA vaccines have been developed against several pathogens, including respiratory syncytial virus (RSV)^18–20^. Following administration, host cells translate the mRNA into antigenic proteins that are subsequently recognized by the immune system. This process activates both CD4+ helper and CD8+ cytotoxic T cells. In parallel, B cells are activated to produce antigen-specific antibodies. Consequently, mRNA vaccines induce both cellular and humoral immune responses, making them attractive candidates for protection agains intracellular pathogens such as *M. tuberculosis*. Several studies have therefore explored the use of mRNA-based vaccines against tuberculosis^21–27^, including one candidate that has entered clinical trials^28,29^.

*Mycobacterium marinum* is a close relative of *M. tuberculosis*, sharing approximately 3,000 orthologous genes^30^. Furthermore, *M. marinum* causes a disease in zebrafish (*Danio rerio*) that resembles human tuberculosis in terms of host-pathogen interactions and granuloma formation, making it a valuable model for studying mycobacterial pathogenesis^31–34^. Using this model, we have previously identified vaccine antigens that confer protection against mycobacterial infection^35–40^. Building on these studies and our current findings, we sought to determine whether protective antigens identified in zebrafish could be translated into a multivalent vaccine strategy. To address this, we designed nucleoside-modified mRNA vaccines encoding four, six, or nine *M. tuberculosis* antigens corresponding to *M. marinum* genes associated with protection against primary infection or reactivation of latent infection in the zebrafish model. We evaluated their immunogenicity and protective efficacy in mice and demonstrated protection against primary *M. tuberculosis* infection comparable to that conferred by BCG vaccination. Furthermore, these vaccines enhanced protection in BCG-vaccinated mice. Finally, the vaccine candidate encoding nine antigens protected guinea pigs against *M. tuberculosis* infection.

## Materials and Methods

### Animals

#### Zebrafish

The wild type zebrafish (*Danio rerio*) AB line was obtained from and maintained by the Tampere Zebrafish Core Facility (Tampere University, Finland). Adult zebrafish were maintained in flow-through systems with a light/dark cycle of 14 h/10 h and fed once a day with suitable granularity of Gemma Micro feed (Planktovie, Marseille, France). Embryos were kept in embryonic medium (5 mM NaCl, 0.17 mM KCl, 0.33 mM CaCl_2_, 0.33 mM MgSO_4_, 0.00001% Methylene Blue) in a 28.5°C incubator until 6 days post fertilization (dpf). For the experiments, adult fish (6-12 months old) and larvae were anesthetized with 0.02% Tricaine (3-aminobenzoic acid ethyl ester, pH 7.0) (Merck, Kenilworth, NJ, USA). The wellbeing of the fish was monitored at least once a day and fish showing symptoms of pain or distress were euthanized with an overdose (0.08%) of the anesthetic Tricaine.

#### Mice

In experiments without *M. tuberculosis* challenge, female C57BL/6JRj mice (4-5 weeks of age) were obtained from Janvier Labs (Le Genest-Saint-Isle, France) and housed in a pathogen-free pre-clinical facility at Tampere University (Tampere, Finland). For *M. tuberculosis* challenge experiments, C57BL/6 mice (6–8 weeks of age) were obtained from Australian BioResources (Moss Vale, Australia) and housed at the Centenary Institute Bioresources animal facility (Sydney, Australia) under Biosafety Level 2 (BSL-2) or Biosafety level 3 (BSL-3) conditions. Mice were maintained on a 12-hour light/dark cycle at 21°C and 45–46% relative humidity in individually ventilated cages, with ad libitum access to standard rodent chow and water. Nesting material, wooden blocks, sterile cardboard, and shelters were provided for environmental enrichment. All procedures were performed after a minimum acclimatization period of one week.

#### Guinea pigs

Dunkin-Hartley outbred guinea pigs (350–400 g) were obtained from Elm Hill Laboratories and housed in a BSL-3 laboratory at the Colorado State University Animal Resources facility. Animals were housed four per cage under controlled temperature (21 ± 2°C) and relative humidity (30 ± 5%) conditions, with a 12-h light/dark cycle and ad libitum access to food and water.

### Ethical statements

Animal experiments were carried out in accordance with the Finnish act on the Protection of Animals Used for Scientific or Educational Purposes (497/2013) and the EU Directive on the Protection of Animals Used for Scientific Purposes (2010/63/EU). Permits for zebrafish maintenance, adult zebrafish experiments, and mouse experiments conducted at Tampere University were obtained from the Animal Experiment Board of Finland (permits: ESAVI/44000/2025, ESAVI/21936/2025, ESAVI/4074/2024). Procedures conducted at the University of Technology Sydney were approved by the Sydney Local Health District Animal Welfare Committee (approval number 2024/039). Procedures involving guinea pigs were conducted at Colorado State University (OLAW D16-00345 (A3572-01)) with Institutional Animal Care and Use Committee approval under protocol 7523. Animal welfare was monitored daily for signs of pain or distress, and body weights of mice and guinea pigs were recorded weekly. Prior to ear tagging, mice received oral carprofen (0.1 mL carprofen per 100 mL drinking water).

### A screen for mycobacterial antigens

24 *M. marinum* antigens were selected for screening based on their differential expression in hypoxic (dormant) and aerated (reactivated) *M. marinum* cultures. The generation of hypoxic culture, subsequent aeration, bacterial mRNA extraction and mRNA sequencing have been described previously^40^. DNA vaccines encoding individual antigens were generated by cloning selected genes or gene fragments into pCMV-GFP vectors (a gift from Connie Cepko; Addgene plasmid #11153; http://n2t.net/addgene:11153; RRID:Addgene_11153), as previously described^35^. Primer sequences and cloned gene fragments are listed in **Table S1**. The protective efficacy of the DNA vaccines against reactivation of mycobacterial infection was evaluated using a dexamethasone-induced reactivation model in zebrafish^39^. Briefly, adult zebrafish were intraperitoneally infected with a low dose (15-40 colony forming units, CFU) of *M. marinum* (ATCC 927 strain) to establish latent infection. Five weeks post-infection, fish were immunized intramuscularly with the DNA vaccine constructs (10-12 µg per fish). Five weeks later, reactivation was induced by administration of dexamethasone-containing feed. Dexamethasone treatment (10 mg per fish per day) continued for four weeks. At the end of the experiment, all remaining fish were euthanized, and *M. marinum* burden was quantified from whole-organ blocks by quantitative PCR (qPCR). Procedures for bacterial preparation, bacterial challenge, organ block collection, DNA extraction and qPCR have been described previously^33,40,41^.

### mRNA vaccine preparation

#### Cloning plasmid templates

The pT7TS-rich plasmid backbone was obtained as a gift from Richard Bélanger (Addgene plasmid #99050; http://n2t.net/addgene:99050; RRID:Addgene_99050). Antigen-coding gene sequences were amplified from genomic DNA of *Mycobacterium tuberculosis* strain H37Rv (ATCC; https://www.atcc.org/) and the *egfp* gene was amplified from the egfp-pcDNA3.1(-) plasmid (GenScript, Dayton, NJ, USA). Primer sequences are listed in **Table S2**. PCR products were purified and cloned into the pT7TS-rich plasmid using standard restriction cloning. Recombinant plasmids were transformed into *Escherichia coli* OneShot™ TOP10 competent cells (Invitrogen, Carlsbad, California, USA) and successful cloning was confirmed by colony PCR and sequencing. For *in vitro* transcription, plasmid DNA was isolated from overnight bacterial cultures using the GenElute™ Endotoxin-Free Plasmid Maxiprep Kit (Merck).

#### In vitro transcription of mRNA

Vaccine mRNA was synthetized by *in vitro* transcription using the plasmid templates. Each plasmid encoding an individual antigen was linearized with XbaI FastDigest restriction enzyme (Thermo Scientific, Waltham, Massachusetts, USA), and mRNA synthesis was performed using the MEGAscript™ T7 Transcription Kit (Invitrogen) according to the manufacturer’s instructions. To generate nucleoside-modified mRNA, 1-methylpseudouridine-5’-triphosphate (m^1^ΨTP) (Trilink, San Diego, California) was used in place of uridine-5’-triphosphate (UTP). A 5’ Cap 1 structure (m^7^G(5’)pppN_1_-) and a poly(A) tail were added using the ScriptCap™ Cap1 Capping system (CellScript, Madison, Wisconsin, USA) and the A-Plus™ Poly(A) Polymerase Tailing Kit (CellScript), respectively. The capped and tailed mRNA was purified with lithium chloride precipitation, resuspended in UltraPure™ water (Invitrogen) and analyzed using the Qubit™ BR RNA Assay Kit (Invitrogen) and capillary electrophoresis (SS total RNA 15 nt Kit, 5200 Fragment Analyzer system, Agilent, California, USA) to assess concentration, integrity and poly(A) tail length. mRNA was stored in -80°C until further use.

#### LNP formulation

Synthetized mRNA was encapsulated into lipid nanoparticles using the LipidLaunch^TM^ LNP-102 Exploration Kit (Cayman Chemical, Ann Arbor, Michigan, USA). mRNAs encoding individual antigens were encapsulated separately to ensure equal concentration of each antigen in the vaccine. For LNP synthesis, SM-102, 1,2-distearoyl-sn-glycero-3-PC, cholesterol and DMG-PEG(2000) were mixed in ethanol at molar ratio of 50:10:38.5:1.5. The lipid solution was combined with aqueous mRNA solution (pH adjusted to 4.0 with Sodium acetate) at an ethanol-to-aqueous ratio of 1:3 and a lipid-to-mRNA ratio of 10:1. To promote self-assembly of the mRNA-LNPs, the lipid solution was rapidly added to the mRNA solution under continuous stirring, and the resulting particles were allowed to assemble for 30 min. The LNPs were dialyzed overnight against a 1000-fold volume of PBS (Gibco, Thermo Fisher Scientific) at +4°C using 10kDa Slide-A-Lyzer™ Dialysis Cassettes (Thermo Fisher Scientific) and then concentrated using the Amicon ultra-4 centrifugal 10kDa filters (Millipore, Burlington, Massachusetts, USA). Encapsulated and free mRNA concentrations were quantified using Qubit BR RNA Kit (Invitrogen). For mRNA release, LNPs were treated with 2% Triton™ X-100 (Sigma-Aldrich, St. Louis, Missouri, USA) at 37°C for 10 min. Finally, the LNPs were stored at +4°C until further use. Vaccines were formulated in PBS by combining equal amounts of each antigen-containing LNPs.

### Evaluating mRNA translation in zebrafish embryos

370 pg of nucleoside modified *egfp* mRNA (synthetized using 1-methylpseudouridine-5’-triphosphate), 370 pg of unmodified *egfp* mRNA (synthetized using regular UTP) or 450 pg of pcDNA3.1 plasmid coding for GFP (GenScript) was microinjected into the cytosol of 1-cell stage zebrafish embryos. Cytosolic microinjections were carried out as previously described^42^. Untreated embryos were used as negative controls in the experiment. GFP expression in live larvae was visualized with Lumar V1.1 fluorescence stereomicroscope (Carl Zeiss MicroImaging GmbH, Göttingen, Germany) 24 hours post injection (hpi).

### Evaluating LNP transfection in S2 cells

*Drosophila melanogaster* S2 cell culture and transfection were performed as previously described^43^ with some modifications. Briefly, S2 cells were cultured in Schneider’s Medium (Merck) supplemented with 10% fetal bovine serum (FBS, Sigma-Aldrich), 100 U/mL penicillin and 100 µg/mL streptomycin at 25°C and 5% CO_2_. For transfection, S2 cells were diluted in fresh Schneider’s Medium, transferred into 24-well plates and left to attach for 1 h. When attached, medium was removed and transfection solution containing medium (without supplements) and either 2 µg of LNP-encapsulated *egfp* mRNA (TUmRNA_GFP_ vaccine) or 2 µg of naked *egfp* mRNA was carefully pipetted on the cells. Medium alone was used as an additional negative control for transfections. After 2 h incubation, Schneider’s Medium with supplements was added and the cells were allowed to grow for 72 h. To visualize GFP in transfected cells, cells were diluted, attached on plates and imaged with IX51 inverted fluorescence microscope and Olympus LUCPlan FL N 20x/0.45 objective (Olympus, Tokyo, Japan).

### Zebrafish immunization studies

Adult male fish were intramuscularly immunized by injecting 5 µl of vaccine or PBS into the dorsal muscle of anesthetized fish using a microinjector (PV830 Pneumatic PicoPump, World Precision Instruments, Sarasota, FL, USA). To visualize GFP expression at the injection site, fish were injected with 0.1 µg of TUmRNA_GFP_ vaccine or PBS and imaged using Nikon AZ100 macroscope (Nikon, Minato City, Tokyo, Japan) at 1, 3 and 5 days post-injection (dpi). To assess the protective efficacy of TUmRNA_6_ in zebrafish, fish were immunized with 0.5 µg of TUmRNA_6_ and boosted three weeks later with either TUmRNA_6_ or PBS. Control fish received two doses of PBS or TUmRNA_GFP_ vaccine with a PBS booster. Three weeks after final dose, fish were challenged with a low dose (∼100 CFU) of *M. marinum* (ATCC 927 strain) via intraperitoneal injection. Bacterial preparation and challenge procedures were performed as previously described^33,41^. Following challenge, fish were monitored at least once a day until sacrifice for the assessment of bacterial burden at 4 weeks post-infection. Bacterial burden was determined from whole organ blocks by quantifying *M. marinum* genome copies using qPCR. Procedures for organ block collection, DNA extraction and qPCR have been previously described^33,41^.

### Mouse vaccination and aerosol infection

Mice were anesthetized with gaseous isoflurane to minimize distress. Mice were vaccinated subcutaneously with 2-5×10⁵ CFU of *Mycobacterium bovis* BCG (100-200 μl in endotoxin-free saline) or intramuscularly in the hind leg with endotoxin-free PBS (50 μl per leg) or one of the mRNA vaccine candidates (50 μl per leg) using insulin syringes. Cultures of *M. tuberculosis* H37Rv were grown in Middlebrook 7H9 broth at 37°C with 5% CO₂ until logarithmic phase was reached (OD₆₀₀ = 0.6–0.8) and frozen at -80°C. Calibrated stocks of bacteria were centrifuged at 3 270 x g for 10 min, resuspended in PBS containing 0.05% Tween-80, then centrifuged at 872 x g for 10 min to pellet bacterial clumps. The suspension was diluted with sterile water to a concentration of 10⁶ CFU/mL and sonicated for 10 s. Four weeks after the final vaccination, mice were exposed to aerosolized *M. tuberculosis* by loading 5.5 mL of bacterial suspension into a Middlebrook airborne infection apparatus (Glas-Col, Terre Haute, IN, USA) to deliver a low-dose infectious inoculum of approximately 100 bacilli per mouse^44^.

### Tissue harvest and single-cell preparation

In the experiments assessing antigen immunogenicity without *M. tuberculosis* challenge, blood samples were collected from the tail vein on days 21 and 43 and centrifuged at 2000 x g for 10 min at +4°C to collect serum. On day 46, mice were euthanized by CO_2_ asphyxiation and bled by cardiac puncture. Serum was isolated using Microtainer SST Blood Collection Tubes (Becton Dickinson, Franklin Lakes, New Jersey, USA) according to the manufacturer’s instructions and stored at -80°C until analysis. Spleen and inguinal lymph nodes were collected into PBS and immediately processed at room temperature. Single-cell suspensions were prepared by mechanical dissociation and filtration through 40 µm cell strainers (Corning, New York, United States) in PBS supplemented with 2% fetal bovine serum (FBS, Sigma-Aldrich) and 2 mM EDTA. Splenocyte suspensions were treated with ACK Lysing Buffer (Gibco) for 1 min to remove erythrocytes. After washing and aggregate removal, resulting cells were resuspended in RPMI 1640 Medium, GlutaMAX™ (Gibco) supplemented with 10% FBS and 1% Penicillin-streptomycin and counted using Via2-Cassette^TM^ (Chemometec, Allerod, Denmark) and NucleoCounter® NC-202^TM^ (Chemometec).

In the *M. tuberculosis* challenge experiments, mice were euthanized at 4 weeks post-infection by CO_2_ asphyxiation. Lungs were perfused with 10 mL of PBS. The right lobe was placed in water for homogenization and bacterial enumeration. The middle-left lobe was perfused with 10% neutral buffered formalin and fixed in 10% neutral-buffered formalin. The remaining lobes were placed in complete RPMI (RPMI 1640 supplemented with 10% FCS, 0.05 mM 2-mercaptoethanol, and 100 U/mL penicillin-streptomycin). Lung lobes were mechanically dissociated using a gentleMACS Dissociator (Miltenyi Biotec, Germany) according to the manufacturer’s mouse lung protocol, then subjected to enzymatic digestion with 13 μg/mL DNase I (Worthington, Lakewood, NJ, USA) and 50 U/mL Collagenase I (Worthington) at 37°C, 5% CO₂ for 30-45 min. Dissociated tissue was passed through a 70 μm nylon cell strainer, and cells were pelleted. The cell pellet was resuspended for 1 min in ACK lysis buffer (150 mM NH₄Cl, 10 mM KHCO₃, 127 μM EDTA, pH 7.4; Sigma-Aldrich) to remove residual red blood cells, before lysis buffer was inactivated with complete RPMI. Cells were pelleted, resuspended in complete RPMI and passed through another 70 μm cell strainer. Mediastinal lymph nodes were passed through a 70 μm nylon sieve and centrifuged to pellet cells before resuspension in complete RPMI. Cells were counted on a Countess 3 Automated cell counter (Thermo Scientific).

### Mouse lung histology

Formalin perfused lung tissue was paraffin-embedded, sectioned at 5 μm, and stained with haematoxylin and eosin (HCE). Stained slides were imaged at 20x magnification using the Zeiss Axio Scan.Z1 slide-scanning microscope and acquired as a mosaic. Histological scoring was performed in a blinded manner by an independent trained pathologist and assessed for the extent of total pathological change, peribronchiolitis, perivasculitis, and alveolitis. Each parameter was scored on a scale of 0 (no apparent change) to 5 (severe changes), and scores were summed to obtain a total lung pathology score.

### Flow cytometry

In the experiments without *M. tuberculosis* challenge, splenocytes and cells from inguinal lymph nodes were processed in parallel for flow cytometry. Cells were centrifuged at 400 x g for 5 min, at room temperature, and resuspended in PBS. For viability assessment, cells were incubated with a fixable viability dye (BD Horizon™ Fixable Viability Stain 780, Waters Biosciences, Milford, MA, US) for 20 min at room temperature in the dark, followed by quenching with buffer (PBS with 2% FBS and 2 mM EDTA). Subsequently, cells were centrifuged (400 x g, 5 min, 4°C). Fc receptors were blocked by incubating cells with Mouse BD Fc Block™ reagent (Waters Biosciences) for 5 min at 4°C. Cells were then stained with fluorochrome-conjugated antibodies in the presence of BD Horizon™ Brilliant Stain Buffer Plus (Waters Biosciences). Detailed information on antibodies used is provided in **Table S3**. Surface staining was carried out for 20 min at 4°C. After staining, cells were washed twice with cold buffer and resuspended in a final acquisition volume. Single-color compensation controls were prepared using Invitrogen™ UltraComp eBeads Compensation Beads (Thermo Fisher Scientific, Waltham, MA, US) or cells, as appropriate. Cells were analyzed with CytoFlex S (Beckman Coulter, Brea, CA, US). Further data analysis was performed using FlowJo™ Software for Windows, versions 10.7.1 and 10.10.0 (Waters Biosciences).

For surface staining of cells isolated from challenged mice, ∼10⁶ lung or lymph node cells were pelleted and incubated with 50 µl Zombie UV fixable viability dye (BioLegend, San Diego, California) in PBS for 30 min. Fc receptors were blocked with anti-CD16/CD32 (Fc block; clone 2.4G2, Waters Biosciences) in buffer (PBS + 2% FCS and 5 mM EDTA) for 30 min at 4°C. Cells were subsequently stained with cocktails of fluorophore-conjugated monoclonal antibodies in buffer for 30 min at 4°C as described in **Table S4**. Samples were acquired on a BD Fortessa X20 flow cytometer (Waters Biosciences) and data were analyzed using FlowJo software version 10.10.6.

For intracellular cytokine staining, antigen-specific restimulation was performed by incubating 2×10⁶ lung cells overnight with *M. tuberculosis* culture filtrate protein (CFP; 1 μg/mL) (NR-14825 was provided by the Biodefense and Emerging Infections Research Resources Repository (BEI Resources, NIAID, NIH)) at 37°C with 5% CO₂, followed by the addition of Brefeldin A (10 μg/mL; Sigma-Aldrich) for the final 4 hours. Following surface staining as described above, cells were permeabilized using the BD Cytofix/Cytoperm Fixation/Permeabilization Kit (Waters Biosciences) per the manufacturer’s instructions, then stained with fluorophore-conjugated antibodies targeting intracellular cytokines for 30 min at 4°C. All stained cells were pelleted and fixed in 10% neutral-buffered formalin.

### Quantification of anti-GFP antibody

Anti-GFP antibodies were detected in sera from vaccinated and control mice using the Anti-Green Fluorescent Protein Antibody (Anti-GFP) ELISA Kit (MyBioSource, San Diego, California, USA). Serum samples were diluted 1:5 or 1:100 in PBS based on their concentration, and the assay was performed according to the manufacturer’s protocol. Each sample was analyzed in technical duplicate. Optical density was measured at 450 nm using a microplate reader (Victor^2^, PerkinElmer, Waltham, Massachusetts, USA), and sample concentrations were determined from a standard curve.

### Preparing peptide libraries for stimulation

Lyophilized peptides (15 amino acids in length, overlapping by 11 amino acids) spanning the full length of each antigen were purchased from GenScript (Nanjing, China). Individual peptides were dissolved in UltraPure^TM^ water (Invitrogen) and dimethyl sulfoxide (DMSO) and pooled to generate antigen-specific peptide libraries at concentrations of 200-400 µg/mL. Peptide libraries were stored at -80°C until use. To generate the peptide pools comprising the six or nine antigens included in the TUmRNA_6_ or TUmRNA_9_ vaccines, respectively, equal amounts of the antigen-specific peptide libraries were combined. For *ex vivo* stimulation, individual peptide libraries and the vaccine-specific peptide pools were diluted in cell culture medium and used at a final concentration of 2 µg/mL.

### Mouse IFNγ ELISpot assay

Antigen-specific T-cell responses were assessed using an enzyme-linked immunospot (ELISpot) assay (Mouse IFNγ single color ELISPOT Kit, Cellular Technology Limited, Ohio, USA) with splenocytes isolated from vaccinated mice. Briefly, splenocytes were resuspended in serum-free CTL-Test^TM^ medium (Cellular Technology Limited) and seeded at 3×10^5^ cells per well in duplicate or triplicate. Cells were stimulated with antigen-specific, GFP-specific, or vaccine-specific peptide pools at 37°C in 5% CO₂ for 16 h. Wells containing medium alone served as negative controls, while cells stimulated with concanavalin A (10 µg/mL; Sigma-Aldrich) and anti-CD28 antibody (0.2 µg/mL; Invitrogen) served as positive controls. Following incubation, the assay was completed according to the kit protocol. Plates were then air-dried for 24 h and scanned using an ImmunoSpot Analyzer (Cellular Technology Limited).

### IFNγ ELISA and multiplex cytokine assay

Mouse splenocytes were seeded onto 24-well plate at 2.5×10^6^ cells per well with 1 mL of RPMI 1640 Medium GlutaMAX™ (Gibco) supplemented with10% FBS and 1% Penicillin-streptomycin. Cells were stimulated with antigen-specific or GFP-specific peptide libraries at 37°C with 5% CO₂ for 68 h. Wells containing medium alone served as negative controls, while cells stimulated with Anti-CD28 (1 µg/mL) and Anti-CD3 (0.5 µg/mL; Invitrogen) antibodies served as a positive control. Following stimulation, the culture supernatants were collected and frozen at−80°C until analysis.

IFNγ, IL1B, IL2, IL4, IL5, IL10, IL13, KC-GRO, IL12p70 and TNF were quantified from cell supernatants using a U-PLEX Th1/Th2 Combo (mouse) Kit (Meso Scale Discovery, Rockville, MD, US) and MESO Quickplex SQ 120 instrument (Meso Scale Discovery) according to the manufacturer’s instructions. Additionally, IFNγ concentrations in the cell supernatants were quantified using an ELISA Pro: Mouse IFNγ Kit (Mabtech, Stockholm, Sweden) according to the Kit protocol.

### Cytometric Bead array

To assess antigen-specific cytokine responses in challenged mice, 1×10⁵ lung cells were plated in 96-well flat-bottom plates in 200 μl of complete RPMI and stimulated for 72 h at 37°C with 5% CO₂ with one of the following: Concanavalin A (ConA; 10 μg/mL), CFP (1 μg/mL), TUmRNA_6_-specific peptide pool (2 μg/mL), or medium. Following stimulation, supernatants were harvested and stored at –80°C until analysis. Concentrations of IFNγ, TNF, IL2, and IL17A were measured using the BD™ Cytometric Bead Array (CBA) kit according to the manufacturer’s instructions. Beads were acquired on a BD Fortessa X20 flow cytometer and data were analyzed using FCAP Array software version 3.0 (BD Biosciences)^45^.

### Guinea pig vaccination and aerosol infection

BCG vaccination of guinea pigs was performed as a single intradermal injection of 1×10^5^ CFU of BCG Danish-1331 strain. mRNA vaccines or PBS were delivered by intramuscular injection, split across both hind limbs. Animals were exposed to *M. tuberculosis* Erdman by aerosol delivery as previously described^46^. Briefly, exposure to *M. tuberculosis* was performed using the Glas-Col whole-body aerosol exposure device. Saline containing 5 mL of 2×10^6^ CFU of *M. tuberculosis* Erdman strain (BEI Resources, #NR-15404) was added to the nebulizer device. Guinea pigs were exposed to the nebulization for 3 min. All procedures involving *M. tuberculosis* were performed in the BSL3 laboratory facility at Colorado State University by trained laboratory staff approved to work under BSL3 conditions and following approved project protocols by the institutional biosafety committee.

### Bacterial burden in guinea pig tissue

Lung, spleen, and mediastinal lymph node were weighed and 9x volume of sterile 1xPBS added to generate a 10% w/v tissue homogenate. Tissues were homogenized in a Precellys Evolution Touch homogenizer at 7200 RPM for 16 s in 15mL CKMix50 homogenizer tubes (Bertin Corp, Rockville, Maryland, USA). Homogenate was plated in 100 μl volumes in a 10-fold serial dilution series on 7H11 agar containing OADC, carbenicillin and amphotericin B. Final *M. tuberculosis* CFU burdens were calculated per tissue weight after 4 weeks of growth.

### Histopathology in guinea pigs

Lung tissue was fixed in 4% paraformaldehyde for 48 h followed by paraffin-embedding, sectioning and mounting of 5 μm sections using standard histology methods. HCE-stained sections were generated using a HistoCore CHROMAX staining system (Leica Biosystems, Illinois, USA) at Colorado State University (RRID:SCR_023562) and scanned to generate whole slide images on a PhenoImager HT pathology imaging system (Akoya Biosciences, Massachusetts, USA). Custom AI-trained algorithms in Visiopharm software (Denmark), trained for detection of tuberculous lesion morphology in the guinea pig model by the Podell laboratory, were used to quantify the total area of tuberculosis lesions and total lung tissue area in vaccinated and unvaccinated guinea pigs. Supervised review by a pathologist was performed prior to final quantification. Lesion burden was calculated as the proportion of detected tuberculous lesion area to total lung tissue area in each HCE-stained section and represented as the mean across three analyzed lung sections from each animal.

### ELISpot for guinea pig cytokine responses

PBMC collected antemortem 1 day prior to *M. tuberculosis* exposure were stimulated with 2 μg/mL of TUmRNA_6_-or TUmRNA_9_-specific peptide pools, or phorbol 12-myristate 13-acetate (PMA) + ionomycin (1x Cell Stimulation Cocktail, BioLegend, San Diego, USA) for positive control stimulation. Cells were left unstimulated as background negative controls for comparison. The ELISpot assays were performed as previously described^47,48^. IFNγ-producing cells were detected with VE4 mouse monoclonal capture antibody and NG3 mouse monoclonal detection antibody. All positive control stimulation conditions, antigen stimulations, and negative controls were performed in duplicate for each animal with 2×10^5^ PBMC per well.

### Statistical analysis

Graphs were generated and the statistical analyses were performed using GraphPad Prism 10. Mann-Whitney test was used to compare two groups with nonparametric distribution whereas Welch’s t-test was used to compare Log_10_ bacterial counts between two groups. For comparisons among three or more groups, one-way ANOVA with Tukey’s multiple comparisons test, one-way ANOVA with Šidák’s multiple-comparisons test, Kruskall-Wallis with Dunn’s multiple comparisons test, or Mann-Whitney with Bonferroni-Holm correction were applied. A p-value of less than 0.05 was considered statistically significant.

## Results

### Identification of candidate antigens for vaccines targeting reactivation of latent mycobacterial infection

We previously established a zebrafish model to identify vaccine antigens that protect against reactivation of mycobacterial infection and demonstrated protective effects of DNA vaccines encoding *M. marinum* counterparts for *M. tuberculosis* genes Rv1886c (encoding Antigen 85B, Ag85B), Rv1009 (encoding Resuscitation-promoting factor B, RpfB) and Rv1234 (encoding uncharacterized protein) in this system^39^. To identify additional *M. marinum* genes associated with the reactivation of latent infection, we performed a differential gene expression analysis of hypoxic (dormant) and reaerated (reactivated) bacterial cultures^40^. That study identified a total of 379 differentially expressed genes, of which 199 genes were upregulated at 2 h or 12 h following reaeration compared with hypoxic or normally growing cultures in logarithmic phase. Seven genes that were upregulated at both 2 h and 12 h after reaeration compared to hypoxic cultures were further analyzed, and their potential as vaccine antigens protecting against reactivation of latent *M. marinum* infection was evaluated in zebrafish^40^. In the present study, we expanded the analysis by selecting 24 genes that possess *M. tuberculosis* homologs and were upregulated either at 2 h or at 12 h post-reaeration compared with hypoxic or normal cultures, or that were exclusively expressed during reactivation (**Table S1**). DNA vaccines encoding each candidate gene were generated and evaluated in the zebrafish reactivation model (**Figure 1A**). Of the 24 genes tested, only MMAR_0678 immunization provided significant protection against reactivation (**Figure 1B** and **Figure S1**). Together with the previously identified *M. marinum* genes exhibiting anti-reactivation properties, MMAR_0678, or its *M. tuberculosis* homolog Rv0359, represents a promising new target for vaccine development against mycobacterial reactivation (**Figure 1C**).

**Figure 1.**
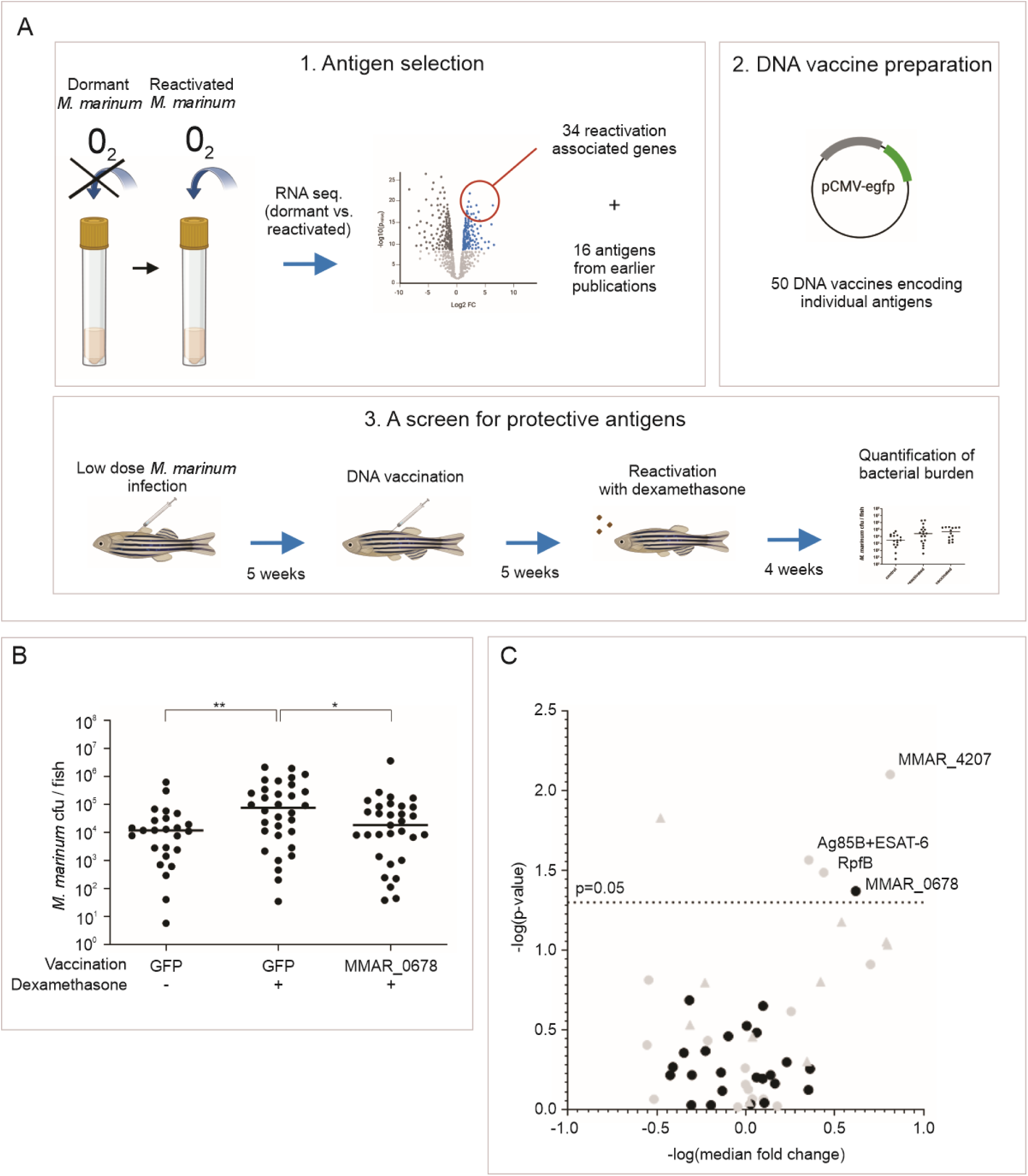
A screen for mycobacterial genes identifies candidate antigens associated with reactivation. **A)** Schematics of the antigen screening strategy. 1) Thirty-four reactivation-associated genes, together with 16 literature-derived antigens, were selected for screening as DNA vaccine candidates. 2) Fifty DNA vaccines encoding antigen-GFP fusion proteins were generated by restriction cloning. 3) Vaccine efficacy was assessed in a zebrafish model: latent infection was established by low-dose intraperitoneal injection, followed by vaccination five weeks post-infection. Five weeks later, reactivation was induced with dexamethasone. Bacterial burden was measured after 4 weeks of treatment. Schematic was created using BioRender (https://www.biorender.com/) and CorelDRAW Graphics Suite 2026 (Corel Corporation, Ottawa, Canada). **B)** Vaccination with a DNA vaccine encoding MMAR_0678 reduced bacterial burden after reactivation compared with the GFP-encoding control vaccine. Data are pooled from two independent experiments (n= 9-18 per group). Dots indicate colony forming units (CFU) in single fish and lines indicate medians. **C)** Volcano plot summarizing the protective effects of the screened *M. marinum* genes. Grey dots denote antigens previously reported by Myllymäki et al. (2018)^39^, grey triangles indicate those described by Niskanen et al. (2020)^40^, and black dots represent unpublished candidates. In **B** and **C**, Statistical significance was determined using the two-tailed Mann-Whitney test. *=p<0.05, **=p<0.01.

### An mRNA vaccine encoding GFP is efficiently translated, well-tolerated and immunogenic in mice

Following a screen that identified potential vaccine antigens against *M. marinum*, we set out to develop a vaccine targeting the human pathogen *M. tuberculosis*. Although antigen discovery was performed using a DNA vaccine platform, advances in mRNA vaccine technology and their demonstrated efficacy and safety in humans^49^ prompted us to transition to an mRNA vaccine platform. We designed a vaccine mRNA encoding a selected antigen, flanked by 5’ and 3’ untranslated regions (UTRs) and containing 5’ cap and 3’ Poly(A) tail (**Figure 2A**). To minimize unwanted and nonspecific innate immune activation, modified uridine nucleotides were incorporated during *in vitro* transcription^50,51^. For delivery, mRNA was encapsulated into lipid nanoparticles (LNPs)^52^.

**Figure 2.**
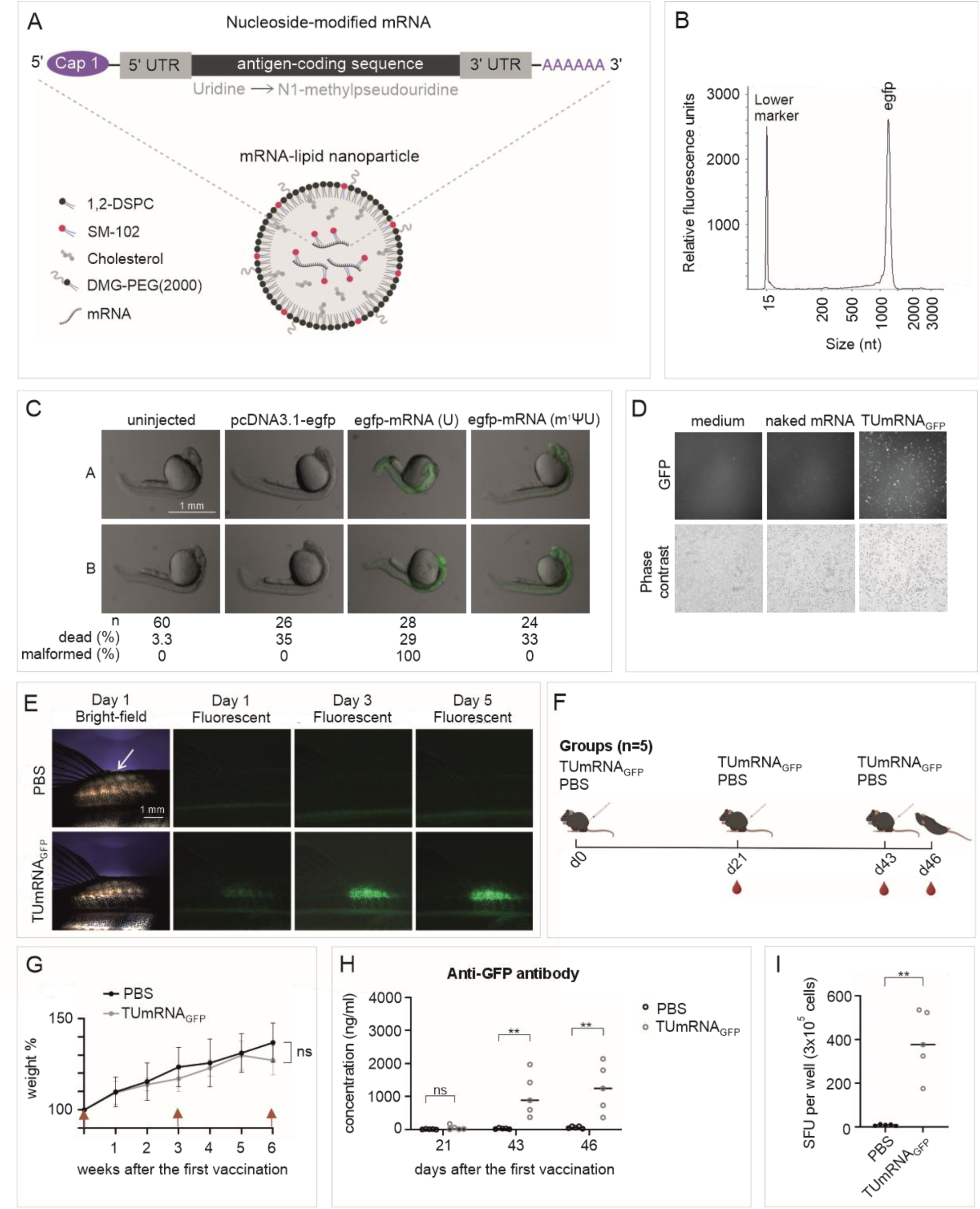
A GFP-encoding mRNA vaccine is well tolerated, efficiently expressed in zebrafish, and elicits antigen-specific immune responses in mice. **A)** Schematic of the mRNA vaccine construct and lipid nanoparticle formulation. The schematic was created using BioRender and CorelDRAW Graphics Suite 2026. **B)** Representative capillary electropherogram of *egfp* mRNA. **C)** GFP expression 24 h after injection of GFP-encoding plasmid DNA, nucleoside-modified mRNA (m¹ΨU) or unmodified mRNA (U). Representative images are shown together with survival and developmental outcomes. **D)** LNP-encapsulated mRNA is efficiently taken up and translated in Drosophila S2 cells. GFP expression was assessed 72 h after transfection of cells with TUmRNA_GFP_ or naked, modified *egfp* mRNA. Representative images are shown. **E)** GFP expression at the injection site (white arrow) following intramuscular administration of TUmRNA_GFP_ in zebrafish. Representative images on days 1, 3, and 5 are shown. In **C-E**, image contrast and/or color were adjusted using ImageJ. **F)** Immunization schedule used to evaluate the immunogenicity of TUmRNA_GFP_ in mice. The schematic was created using BioRender and CorelDRAW Graphics Suite 2026. **G)** Body weight of mice immunized with TUmRNA_GFP_ or PBS expressed as a percentage of the baseline weight (mean+SD, n=5). Red arrows indicate vaccinations. **H)** Anti-GFP antibody in the sera of TUmRNA_GFP_ vaccinated mice. Circles represent antibody concentration in individual mice and lines indicate medians (n=5). **I)** IFNγ ELISpot responses of splenocytes following restimulation with GFP peptide pool. Circles represent mean spot-forming units (SFU) per 3×10^5^ cells from duplicate wells and lines indicate medians (n=5). In **G-I**, statistical significance was determined using the Mann-Whitney test, **=p<0.01.

Before evaluating zebrafish-derived tuberculosis antigens, we first established that our mRNA platform supported antigen expression and immunogenicity across multiple model systems by generating and testing an mRNA vaccine encoding green fluorescent protein (GFP). The quality of the vaccine mRNA, including correct size, successful Poly(A) tail synthesis, and high integrity, was verified by capillary electrophoresis (**Figure 2B**). Translation and tolerability of the vaccine mRNA was then assessed by injecting modified *egfp* mRNA into the cytosol of zebrafish embryos at 1-cell stage and monitoring early development and GFP expression at 24 hours post injection (hpi). For comparison, embryos were injected with either unmodified *egfp* mRNA or GFP-encoding plasmid (pcDNA3.1-egfp). Robust GFP expression was detected 24 hpi in embryos injected with both modified and unmodified mRNA, whereas no GFP was detected in embryos injected with the egfp-encoding plasmid (**Figure 2C**). Survival rates were comparable across all groups and no developmental defects were observed in embryos injected with plasmid or modified mRNA. In contrast, 100 % of embryos injected with unmodified mRNA exhibited clear developmental abnormalities, including curved tails. These results demonstrate efficient translation of the modified mRNA and improved tolerability relative to unmodified mRNA.

Next, we evaluated the transfection efficiency of our LNP-encapsulated mRNA *in vitro* and *in vivo* using a GFP-encoding mRNA vaccine (TUmRNA_GFP_). Incubation of S2 cells with TUmRNA_GFP_ resulted in robust GFP expression, indicating successful and efficient delivery and translation of the vaccine mRNA (**Figure 2D**). No GFP expression was detected following exposure to naked *egfp* mRNA or medium alone. Transfection and translation of TUmRNA_GFP_ were further assessed in adult zebrafish. Following intramuscular injection, GFP expression was readily detected over 5 days post injection (dpi), confirming efficient *in vivo* expression of the encoded antigen (**Figure 2E**).

Having shown the tolerability and efficient antigen expression in zebrafish, we next examined the immunogenicity of TUmRNA_GFP_ in mice. Female C57BL/6JRj mice were intramuscularly immunized with 2 µg of TUmRNA_GFP_ or PBS, three times, at three-week intervals (**Figure 2F**). Throughout the experiment, mouse weight and general well-being were monitored, and no adverse effects or statistically significant differences in weight were observed between the groups (**Figure 2G**).

Measurement of antigen-specific immune responses demonstrated that anti-GFP antibodies became detectable in the serum of TUmRNA_GFP_-vaccinated mice after the second immunization (**Figure 2H**). To assess cellular immune responses induced by TUmRNA_GFP_, splenocytes from vaccinated and control mice were stimulated on IFNγ ELISpot plates with a GFP peptide library. Splenocytes isolated from TUmRNA_GFP_-vaccinated mice generated significantly higher numbers of IFNγ-secreting cells than those from PBS-treated controls (**Figure 2I**). Collectively, these data demonstrate that TUmRNA_GFP_ is well tolerated, supports efficient *in vivo* antigen expression, and is immunogenic in mice, eliciting both antigen-specific humoral and cellular immune responses.

### TUmRNA_6_ protects zebrafish against *M. marinum* infection and induces antigen-specific memory T-cell response in mice

Our antigen screen identified a total of nine *M. tuberculosis* antigens whose *M. marinum* homologs conferred protection in the zebrafish model of tuberculosis. These included four antigens, PE15, PE13, RpfE and Cdh, that protected against primary *M. marinum* infection^37^, three antigens, Rv0359, Rv1234 and RpfB, that protected against reactivation, and two antigens, Ag85B and ESAT-6, that conferred protection in both models^35,39^ (**Table 1**). Based on these findings, we designed a multivalent mRNA vaccine encoding six antigens: Ag85B, RpfE, PE15, PE13, Rv1234 and Rv0359. As for TUmRNA_GFP_, the size and integrity of each vaccine mRNA were verified by capillary electrophoresis (**Figure 3A**). Then, individual mRNAs were encapsulated into LNPs and all six were combined in equal amounts to generate the multivalent TUmRNA_6_ vaccine.

**Figure 3.**
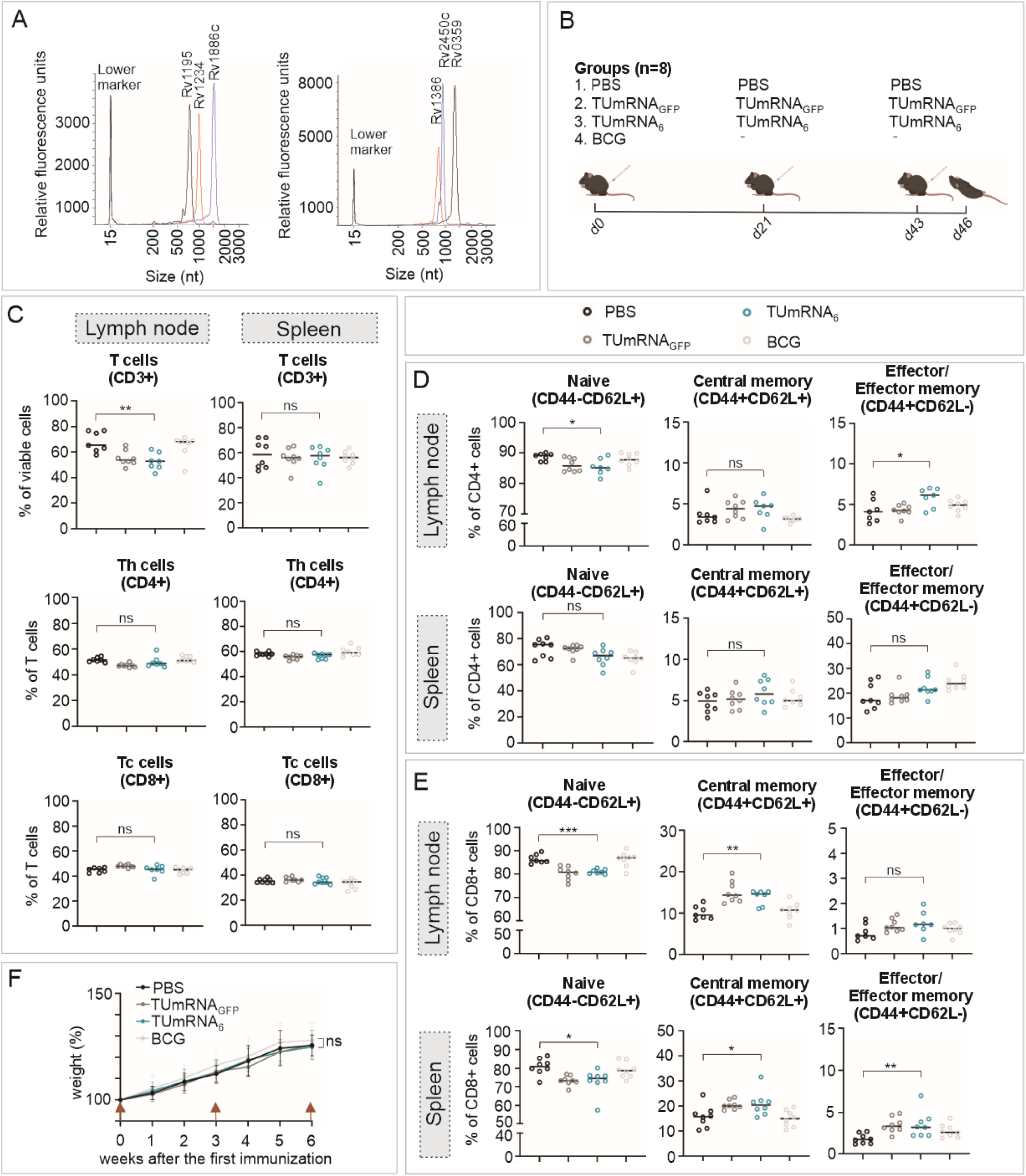
TUmRNA_6_ vaccination promotes memory T cell differentiation in mice. **A)** Representative capillary electropherograms of the mRNAs encoding the six TUmRNA_6_ antigens. **B)** Immunization schedule used to assess the immunogenicity of TUmRNA_6_ in mice. The schematic was created using BioRender and CorelDRAW Graphics Suite 2026. **C)** Frequencies of total CD3+ T cells and CD4+ and CD8+ T-cell subsets in spleens and inguinal lymph nodes. **D)** Frequencies of naive (CD44−CD62L+), central memory (CD44+CD62L+), and effector memory (CD44+CD62L−) CD4+ T cells. **E)** Frequencies of naive, central memory, and effector memory CD8+ T cells. In **C-E**, circles represent individual mice, and lines depict medians (n=7-8 per group). **F)** Body weight of mice immunized with PBS, TUmRNA_GFP_, TUmRNA_6_, or BCG, expressed as percentage of baseline weight (mean + SD, n=13-14, one BCG-vaccinated mouse was excluded for health issues unrelated to vaccination); red arrows indicate vaccinations. Data were pooled from three experiments. In **C-F**, statistical significance was determined using the Mann–Whitney test, *=p<0.05, **=p<0.01, ***p<0.001ns=not significant.

**Table 1.** Vaccine antigens identified in zebrafish. The protective effects of *M. marinum* genes were tested in zebrafish as DNA vaccines. Corresponding *M. tuberculosis* genes were used as vaccine antigens in mRNA vaccine candidates. *a DNA vaccine encoding three antigens: MMAR_2777, MMAR_0188 and MMAR_5449. **a DNA vaccine encoding two antigens: MMAR_2777 and MMAR_0188.

| *M. marinum* antigens identified in zebrafish |  |  |  | Corresponding *M. tuberculosis* antigens |  |  |  |
| Gene | Protective against | Protection in zebrafish | Reference | Gene/protein (% homology) | Vaccine candidate |  |  |
|---|---|---|---|---|---|---|---|
| MMAR_5258 | primary infection | 56% (p<0.01) | Myllymäki et al. 2017<sup>37</sup> | Rv1386/PE15 (67%) | TUmRNA<sub>4</sub> | TUmRNA<sub>6</sub> | TUmRNA<sub>9</sub> |
| MMAR_4241 | primary infection | 50% (p<0.05) | Myllymäki et al. 2017<sup>37</sup> | Rv1195/PE13 (70%) |  |  |  |
| MMAR_3776 | primary infection | 88% (p<0.05) | Myllymäki et al. 2017<sup>37</sup> | Rv2450c/RpfE (74%) |  |  |  |
| MMAR_2777 | primary infection * | 64% (p<0.03) | Oksanen et al. 2013<sup>35</sup> | Rv1886c/Ag85B (89%) |  |  |  |
|  | reactivation** | 56% (p=0.02 | Myllymäki et al. 2018<sup>39</sup> |  |  |  |  |
| MMAR_0678 | reactivation | 76% (p=0.02) |  | Rv0359 (70%) |  |  |  |
| MMAR_4207 | reactivation | 85% (p=0.003) | Myllymäki et al. 2018<sup>39</sup> | Rv1234 (95%) |  |  |  |
| MMAR_4479 | reactivation | 63% (p=0.01) | Myllymäki et al. 2018<sup>39</sup> | Rv1009/RpfB (85%) |  |  |  |
| MMAR_3445 | primary infection | 62% (p<0.05) | Myllymäki et al. 2017<sup>37</sup> | Rv2289/Cdh (68%) |  |  |  |
| MMAR_0188 | primary infection* | 64% (p<0.03) | Oksanen et al. 2013<sup>35</sup> | Rv3875/ESAT-6 (90%) |  |  |  |
|  | reactivation** | 56% (p=0.02) | Myllymäki et al. 2018<sup>39</sup> |  |  |  |  |

To assess vaccine tolerability and protection against related mycobacteria, adult zebrafish were vaccinated once or twice with TUmRNA_6_ and infected with approximately 100 CFU of *M. marinum* three weeks later (**Figure S2**). Fish were monitored daily, and no adverse effects were detected following vaccination. Bacterial burden was significantly reduced in TUmRNA_6_-vaccinated fish compared with PBS-treated controls after four weeks. Median bacterial counts were 2.1×10⁴ and 2.8×10⁴ in fish receiving one or two TUmRNA_6_ vaccinations, respectively, compared with 1.7×10⁵ in PBS-treated fish (p=0.02) (**Figure S2**). These findings demonstrate that TUmRNA_6_ protects against *M. marinum* infection, and like BCG, may provide cross-protection against related mycobacterial species.

Next, we evaluated immunogenicity of TUmRNA_6_ in mice. To gain insight into optimal vaccine dose, female C57BL/6JRj mice were immunized three times at three-week intervals with either a low dose (0.33 µg per antigen; 2 µg total) or a high dose (2 µg per antigen; 12 µg total) of TUmRNA_6_. PBS-immunized mice served as controls. Although mice receiving the high dose exhibited a slight reduction in weight gain following the second immunization compared with PBS-treated controls, both vaccine doses were well tolerated, with no adverse effects observed during follow-up (**Figure S3**). Three days after the final immunization, splenocytes were harvested and restimulated with a TUmRNA_6_-specific peptide pool. TUmRNA_6_ vaccination elicited robust antigen-specific IFNγ responses, with significantly higher numbers of IFNγ-secreting splenocytes in vaccinated mice than in PBS-treated controls (**Figure S3**). IFNγ responses were comparable between the low- and high-dose groups. As increasing the vaccine dose did not significantly enhance the cellular immune response, the low-dose regimen was selected for subsequent mouse studies.

To further evaluate the immunogenicity of TUmRNA_6_, female C57BL/6JRj mice were immunized three times with 2 µg of TUmRNA_6_, 2 µg of TUmRNA_GFP_ or PBS, or once with BCG. Three days after the final immunization, mice were euthanized and spleen and inguinal lymph node cell populations were assessed by flow cytometry (**Figure 3B**). The gating strategies are shown in **Figure S4**. Overall, TUmRNA_6_-vaccination was associated with a shift from naive to memory T-cells. While the proportion of CD4+ and CD8+ T cells remained unchanged between groups (**Figure 3C**), TUmRNA_6_ vaccination resulted in a significant decrease in naive T-cell subsets and an increase in central memory and effector T cells in both the spleen and lymph nodes (**Figures 3D–E**). Analysis of additional T cell activation markers CD25, CD69, and PD-1 on CD4+ and CD8+ cells demonstrated that only the proportion of CD69+ CD8+ cells was significantly increased in the lymph node of TUmRNA_6_-vaccinated mice compared to the PBS-treated mice (**Figure S5**). TUmRNA_6_-vaccination also affected B cell populations: the proportion of B cells (B220+) in lymph nodes was significantly decreased in vaccinated mice as was the proportion of IgM+ B cells in the spleen (**Figure S5**).

Throughout the experiment, animal well-being and body weight were closely monitored. No adverse effects were observed following immunization, as supported by comparable relative weight changes across all experimental groups (**Figure 3F**). Of note, one TUmRNA_6_-vaccinated mouse exhibited markedly enlarged spleen and lymph nodes at study termination. Collectively, these findings indicate that TUmRNA_6_ is well tolerated and induces the expansion of memory T-cell populations.

### TUmRNA_6_ antigens are immunogenic in mice and induce secretion of multiple cytokines, including IFNγ, by splenocytes

To evaluate the immunogenicity of the six TUmRNA_6_ vaccine antigens individually, splenocytes from vaccinated and control mice were restimulated with peptide pools corresponding to individual antigens or GFP, or with medium alone, and analyzed for IFNγ-producing cells. Antigen-specific IFNγ responses were detected against all six vaccine antigens following restimulation with the corresponding peptide pools and were significantly higher than those observed in PBS-treated and TUmRNA_GFP_-vaccinated control mice (**Figure 4A**). plenocytes from BCG-vaccinated mice showed elevated responses to Ag85B, and PE15. Consistent with our previous observations, splenocytes from TUmRNA_GFP_-vaccinated mice showed a strong response to GFP peptide stimulation. Altogether, these results demonstrate that all six TUmRNA_6_ antigens are immunogenic and induce IFNγ-positive T cell responses in mice.

**Figure 4.**
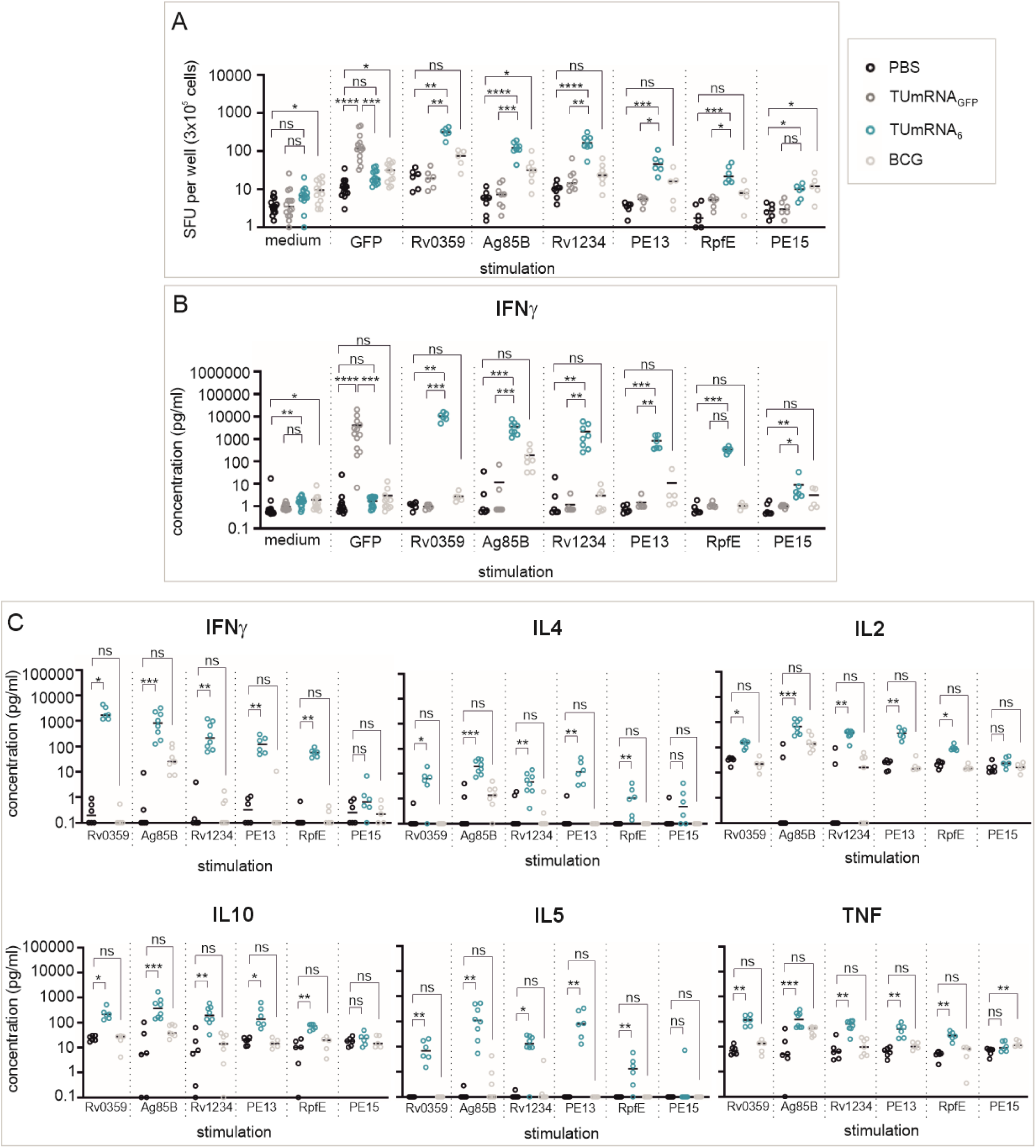
TUmRNA_6_ vaccination elicits antigen-specific cytokine responses in mice. **A)** IFNγ-producing cells were measured by ELISpot after 16 h restimulation of splenocytes with the indicated peptide pools or medium. Data were pooled from three independent experiments. Circles represent mean spot-forming units (SFU) per 3×10^5^ cells from duplicate or triplicate wells and lines indicate medians (n=5–14). **B)** IFNγ concentrations in splenocyte culture supernatants after 68 h restimulation with the indicated peptide pools or medium, measured by ELISA. Circles represent mean concentrations from duplicate wells and lines indicate medians (n=5–14). **C)** Cytokine concentrations in splenocyte culture supernatants after 68 h restimulation with the indicated peptide pools, measured by multiplex cytokine assay. Shown are the six cytokines exhibiting the strongest responses following vaccination. Circles represent individual measurements and lines indicate medians (n=5-8). In **A–C**, one BCG-vaccinated mouse was excluded for health issues unrelated to vaccination. Statistical significance was determined using the Kruskal–Wallis test with Dunn’s multiple-comparisons test; *p<0.05, **p<0.01, ***p<0.001, ns=not significant.

To further examine the response to the vaccine antigens, splenocytes were stimulated with vaccine antigens, GFP or medium alone, and IFNγ in culture supernatants were quantified by ELISA. Consistent with the ELISpot results, splenocytes from TUmRNA_6_-vaccinated mice produced significantly more IFNγ following antigen stimulation compared to PBS-treated or TUmRNA_GFP_-vaccinated controls (**Figure 4B**).

To further characterize the immune response induced by TUmRNA_6_, we analyzed antigen-specific production of multiple Th1- and Th2-associated cytokines using a multiplex assay. Among these, IFNγ, IL2, IL4, IL5, IL10, and TNF were the most highly expressed in splenocytes from TUmRNA_6_-vaccinated mice compared with PBS-treated controls (**Figure 4C**). These responses were most pronounced following stimulation with peptide pools containing Ag85B, Rv1234, Rv0359, PE13, or RpfE whereas responses to PE15 stimulation were more modest and did not reach statistical significance. Splenocytes from BCG-vaccinated mice also responded to stimulation with Ag85B and PE15. Finally, Ag85B stimulation induced a broader cytokine profile, triggering IL13 and IL1B production by splenocytes from TUmRNA_6_-vaccinated mice (**Figure S6**). Supporting assay specificity, splenocytes from TUmRNA_GFP_-vaccinated mice showed a strong response to GFP peptide stimulation, while no increased cytokine production was detected in any group following stimulation with medium alone (**Figure S6**). Taken together, these findings demonstrate that TUmRNA_6_ elicits a broad antigen-specific immune response in mice, encompassing both Th1- and Th2-associated immune pathways, including robust IFNγ production.

### TUmRNA_4_ and TUmRNA_6_ protect mice from *M. tuberculosis* infection

Next, we evaluated whether our vaccine candidates have protective effect in the mouse primary tuberculosis challenge model. We used two vaccine formulations: TUmRNA_6_, comprising six antigens and designed to protect against both primary infection and reactivation of latent infection, and TUmRNA_4_, comprising four antigens (Ag85B, RpfE, PE13, and PE15) and designed primarily to protect against primary infection (**Table 1**). Both vaccines contained a total of 2 µg of mRNA; TUmRNA_6_ included 0.33 µg of each antigen-encoding mRNA, whereas TUmRNA_4_ included 0.5 µg of each mRNA.

Female C57BL/6JRj mice were immunized three times, at three-week intervals, with either 2 µg of TUmRNA_6_, 2 µg of TUmRNA_4_, 2 µg of TUmRNA_GFP_ or PBS (**Figure 5A**). BCG vaccination administered at day 0 was used as a comparator. Four weeks after the final vaccination, mice were infected via aerosol with approximately 100 CFU of *M. tuberculosis*. Bacterial burdens in the lungs and spleen were quantified four weeks post-infection. Pulmonary bacterial loads were significantly lower in mice vaccinated with BCG, TUmRNA_6_ and TUmRNA_4_ compared to PBS or TUmRNA_GFP_-vaccinated controls (**Figure 5B**). Compared to PBS controls, the corresponding average log_10_ reductions in CFU were 1.3, 1.1 and 1.1 for BCG, TUmRNA_6_ and TUmRNA_4_, respectively (p<0.0001 for all, no statistically significant differences between the vaccinated groups). No statistically significant differences in the splenic bacterial loads were observed for individual vaccine groups. However, there was a trend toward reduced CFU in the spleens of TUmRNA_6_- and TUmRNA_4_-vaccinated mice compared to PBS-treated controls. When combining the two mRNA vaccine groups and comparing them with the PBS-controls, mean bacterial counts in the spleen were significantly lower, by 0.7 log_10_, in mRNA-vaccinated mice (p=0.04).

**Figure 5.**
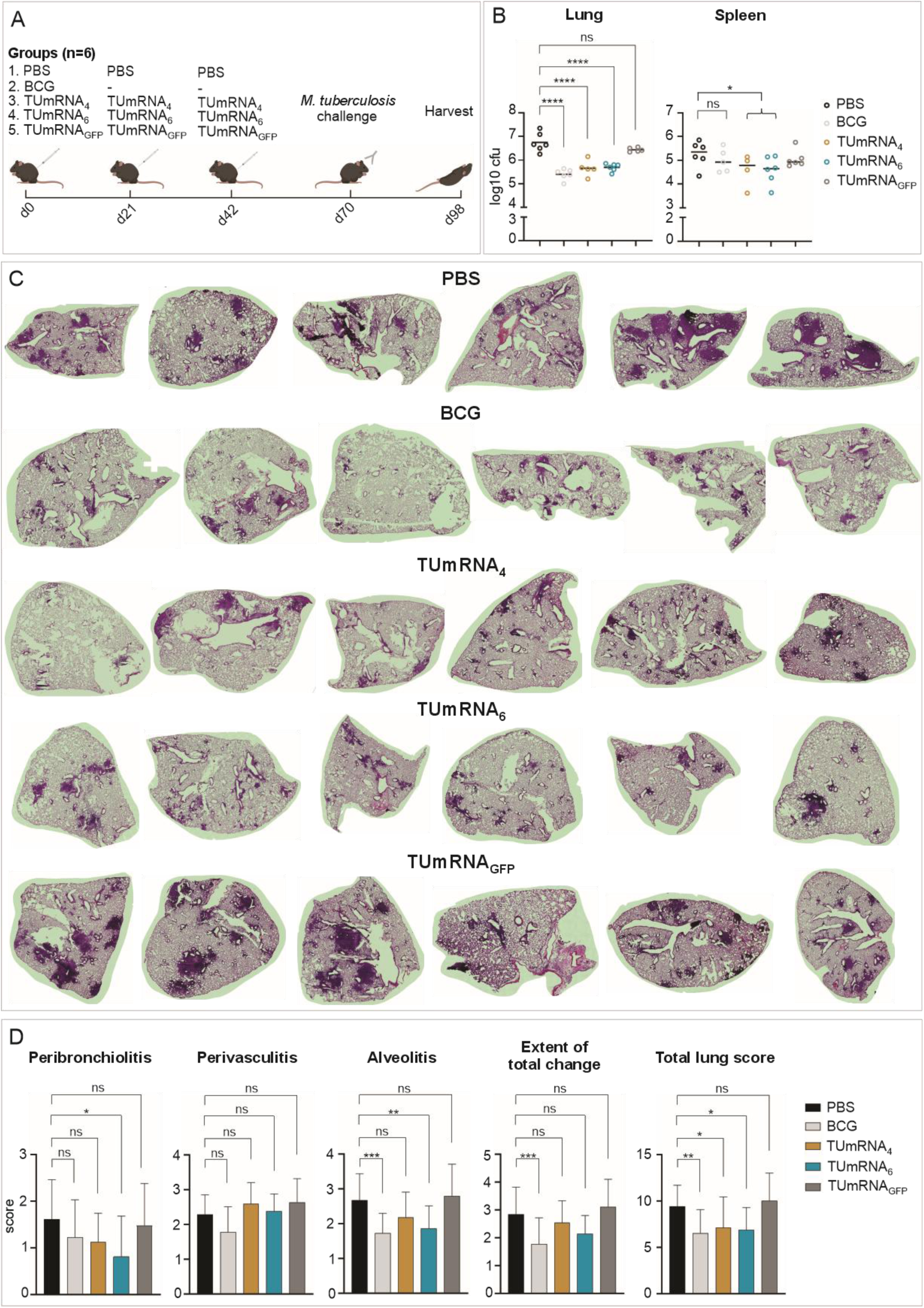
TUmRNA_4_ and TUmRNA_6_ protect against *M. tuberculosis* infection in mice. **A)** Immunization and challenge schedule used to evaluate the protective efficacy of TUmRNA_4_ and TUmRNA_6_ in mice. The schematic was generated using BioRender and CorelDRAW Graphics Suite 2026. **B)** *M. tuberculosis* burden in lungs and spleens of vaccinated mice four weeks after aerosol challenge. Circles represent the log_10_ bacterial counts of individual mice and lines depict group means (n=4-6). **C)** Representative HCE-stained lung sections from vaccinated and challenged mice. **D)** Lung histopathology scores for peribronchiolitis, perivasculitis, alveolitis and overall pathological changes, summed to obtain the total lung score. In **B** and **D**, statistical significance was determined using one-way ANOVA followed by Tukey’s multiple-comparisons test. *=p<0.05, **=p<0.01, ***p<0.001, ****p<0.0001, ns=not significant.

The protective effect of the candidate vaccines was further assessed by lung histology. HCE-stained lung sections revealed significantly reduced inflammation in mice vaccinated with BCG, TUmRNA_6_, or TUmRNA_4_ compared to PBS controls, as reflected by total clinical scores (**Figures 5C-D**). When analyzing individual scoring parameters, peribronchiolitis alveolitis and perivasculitis, TUmRNA_6_ vaccination resulted in significantly decreased scores compared to PBS for the first two parameters. Furthermore, the TUmRNA_6_ showed a trend toward improved histopathological outcomes compared with TUmRNA_4_, and was therefore selected for subsequent studies. Taken together, these results demonstrate that both TUmRNA_6_ and TUmRNA_4_ confer protection against *M. tuberculosis* infection in mice, providing protection comparable to BCG vaccination.

### Two doses of TUmRNA_6_ or TUmRNA_G_ are sufficient for protection against *M. tuberculosis* infection in mice

Having demonstrated the protective efficacy of the four-and six-antigen mRNA vaccines in mice, we next developed a broader multivalent vaccine, TUmRNA_9_, encoding all nine identified antigens. In addition to TUmRNA_6_ antigens, TUmRNA_9_ codes for RpfB, ESAT-6 and Cdh (**Table 1**). Like the TUmRNA_6_ antigens, the three additional antigens in TUmRNA_9_ were immunogenic in mice, as demonstrated by antigen-specific IFNγ production by splenocytes following restimulation with individual peptide pools (**Figure S7**).

To compare the protective effects of TUmRNA_6_ and TUmRNA_9_, mice were immunized either two or three times with either vaccine, infected four weeks after the final vaccination and evaluated for bacterial burden in the lungs and spleens five weeks post-infection (**Figure 6A**). To examine potential sex-specific effects, each group included four male and four female C57BL/6JRj mice. Combined analysis of all mice showed that two- and three-dose regimens with TUmRNA_6_ conferred comparable protection in the lungs, with mean log_10_ reductions in CFU of 0.8 and 0.7, respectively, compared to PBS controls (**Figure 6B**) (p=0.003 and p=0.006). BCG vaccination provided similar protection (0.8 log₁₀ reduction in CFU; p=0.003). TUmRNA_9_ vaccination also provided some protection, however, this reached statistical significance only when the results from the two- and three-dose regimens of TUmRNA_9_ were combined (0.5 log_10_ reduction, p=0.04). A significant reduction in splenic bacterial burden was also observed in mice receiving BCG or two doses of TUmRNA_6_, with log_10_ reductions of 0.9 in CFU relative to PBS-treated mice (p=0.01 and p=0.02, respectively). Decreased bacterial burden in TUmRNA_6_- and TUmRNA_9_-vaccinated mice was also accompanied by a trend toward milder pulmonary inflammation, as assessed by histopathological scoring of HCE-stained lung section. However, these differences did not reach statistical significance relative to PBS-treated mice (**Figure S8**).

**Figure 6.**
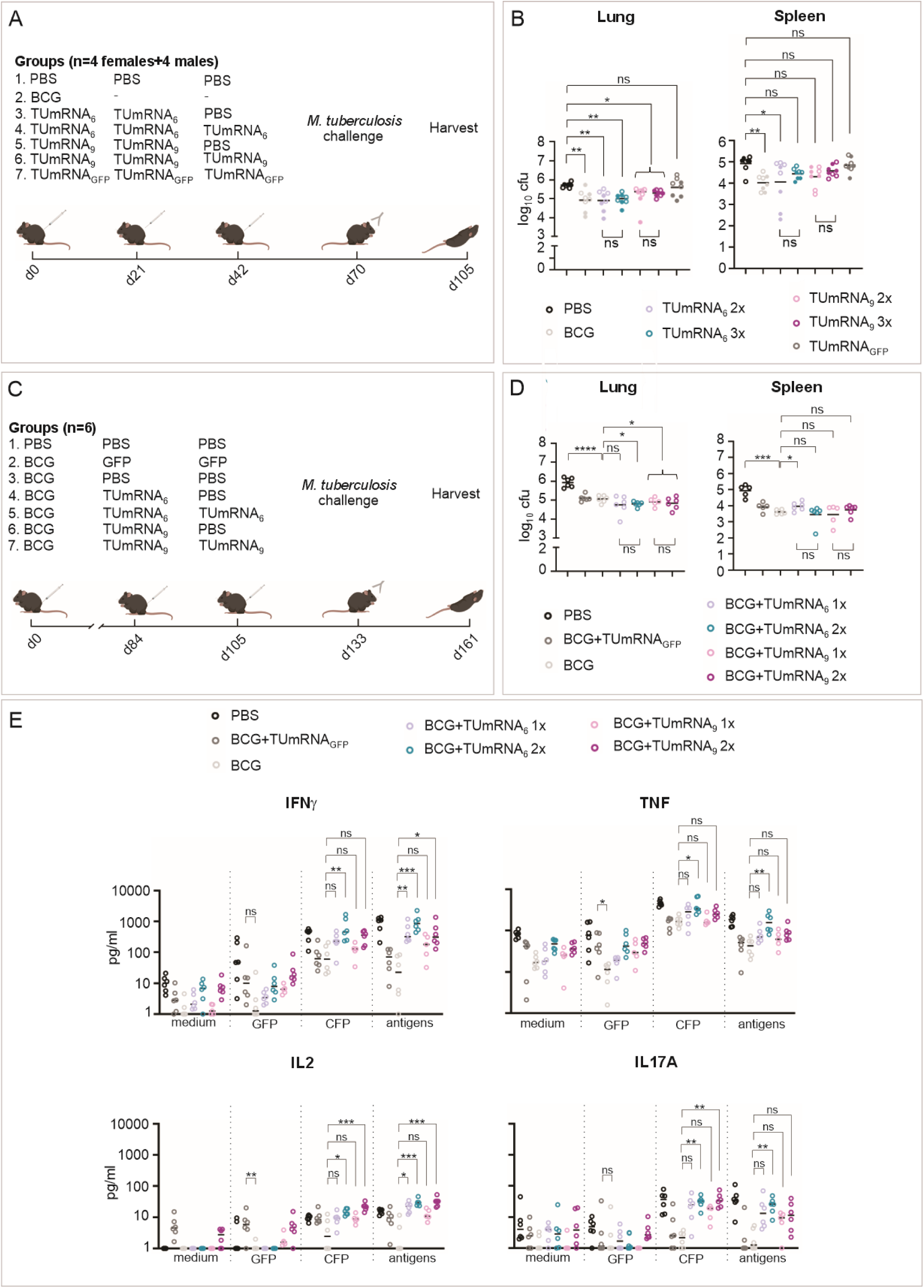
Multivalent mRNA vaccines protect against *M. tuberculosis* and enhance BCG-mediated immunity in mice. **A)** Immunization and challenge schedule used to compare two-dose (2x) and three-dose (3x) regimens of TUmRNA_6_ and TUmRNA_9_. The schematic was generated using BioRender and CorelDRAW Graphics Suite 2026. **B)** *M. tuberculosis* burden in the lungs and spleens of vaccinated and challenged mice. Circles and dots represent the log_10_ bacterial counts in female and male mice, respectively, and lines indicate means (n=8). Statistical significance was determined using one-way ANOVA followed by Tukey’s multiple-comparisons test. *=p<0.05, **=p<0.01, ns=not significant. **C)** Immunization and challenge schedule used to evaluate TUmRNA_6_ and TUmRNA_9_ as booster vaccines following BCG priming. The schematic was generated using BioRender and CorelDRAW Graphics Suite 2026. **D)** *M. tuberculosis* burden in the lungs and spleens of BCG-primed, TUmRNA-boosted and aerosol-challenged mice. Circles represent the log_10_ bacterial counts in individual mice and lines indicate means (n=5-6, one TUmRNA_9_ 1x mouse was excluded from the study due to an injury unrelated to vaccination). Statistical significance was determined using Welch’s t-test. *=p<0.05, ***=p<0.001, ****=p<0.0001, ns=not significant. **E)** Antigen*-*specific cytokine response in the lungs of BCG-primed and boosted mice following restimulation with medium alone, GFP peptide pool, CFP or a TUmRNA_6_-specific peptide pool, measured by cytometric bead array. Circles indicate mean values from duplicate measurements, and lines represent group medians (n=5-6, one TUmRNA_9_ 1x mouse was excluded from the study due to an injury unrelated to vaccination). Values below 1 were assigned a value of 1 for visualization. Statistical significance was determined using the Kruskal-Wallis test with Dunn’s multiple-comparisons test. *=p<0.05, **=p<0.01, ***=p<0.001, ns=not significant.

To characterize the cellular immune responses induced by TUmRNA_6_ and TUmRNA_9_, lung and lymph node cells from vaccinated and control mice were analyzed by flow cytometry (gating strategy shown in **Figure SG**). Vaccination induced phenotypic changes consistent with memory T-cell differentiation. In the lungs, TUmRNA_6_ or TUmRNA_9_ significantly increased the frequency of central memory T cells and decreased the frequency of effector memory T cells compared with PBS-treated controls (**Figure S10**). In the lymph nodes, TUmRNA_6_- and TUmRNA_9_-vaccinated mice exhibited lower frequencies of KLRG1+ T cells and short-lived effector cells (SLECs; KLRG1+CD127−) (**Figure S11**). Overall, the patterns were generally consistent following both two- and three-dose vaccination regimens.

These data show that two immunizations are sufficient to induce protective immunity against primary *M. tuberculosis* infection in mice. The inclusion of three additional antigens in TUmRNA_9_ did not result in improved protection relative to TUmRNA_6_ in this model.

### Multivalent mRNA vaccines enhance BCG-induced immune response and protection against *M. tuberculosis* in mice

Given the widespread use of BCG in infant immunization programs worldwide, novel tuberculosis vaccine candidates should be evaluated for their safety and efficacy as booster vaccines following BCG priming. To investigate this, female C57BL/6JRj mice were primed with BCG and, 12 weeks later, received one or two booster doses of either TUmRNA_6_ or TUmRNA_9_. Four weeks after the final booster vaccination, mice were challenged with *M. tuberculosis* and bacterial burden and antigen-specific immune responses were assessed four weeks post-infection (**Figure 6C**).

As expected, BCG-primed mice receiving either PBS or TUmRNA_GFP_ as a control booster had significantly lower pulmonary bacterial burdens than unvaccinated PBS-treated controls, with mean reductions of 0.9 and 0.8 log_10_ CFU, respectively (p<0.0001 and p=0.0001) (**Figure 6D**). Boosting with TUmRNA_6_ further reduced pulmonary bacterial burden relative to BCG alone, with both one-and two-booster regimens resulting in an additional 0.3 log_10_ reduction in mean CFU compared with BCG alone. However, owing to the higher variability observed in mice receiving a single TUmRNA_6_ booster, statistical significance was reached only in the group receiving two booster doses (p=0.02). Similarly, one or two TUmRNA_9_ booster doses resulted in an additional 0.2 log_10_ reduction in mean pulmonary bacterial burden compared with BCG alone, although neither regimen individually reached statistical significance. However, when both vaccine regimens were analyzed together, pulmonary bacterial burden was significantly reduced compared with the BCG-primed, PBS-boosted control group (p=0.04).

In a separate experiment assessing longer-term protection, mice were challenged four weeks after the final immunization and bacterial burden was determined 20 weeks post-challenge (**Figure S12**). At this later time point, neither BCG alone nor BCG followed by TUmRNA_6_ or TUmRNA_9_ boosting provided significant protection based on bacterial burden in the lungs or spleen (**Figure S12**). Although protection was not maintained at 20 weeks post-challenge, these findings are consistent with previous reports describing waning BCG-mediated protection in murine models^22,53^.

We next assessed antigen-specific immune responses in BCG-primed and TUmRNA_6_- or TUmRNA_9_-boosted mice following *M. tuberculosis* challenge. Lung cells were cultured with *M. tuberculosis* culture filtrate proteins (CFP) or the TUmRNA_6_ peptide pool, or controls (GFP-peptide pool or medium). Compared with BCG-primed, PBS-boosted mice, TUmRNA_6_ boosting significantly increased antigen-specific IFNγ and IL2 production regardless of whether one or two booster doses were administered (**Figure 6E** and **Figure S13**). Antigen-specific TNF and IL17A productions also significantly increased, although only after administration of two TUmRNA_6_ booster doses. Similarly, two booster doses of TUmRNA_9_ induced antigen-specific IFNγ and IL2 production by the lung cells. However, the frequencies of polyfunctional T cells remained low after TUmRNA boosting, with a slight increase in the IFNγ+IL2+ and IFNγ+TNF+IL2+ CD8+ cells in TUmRNA_9_-boosted mice as measured by intracellular cytokine staining (**Figure S14**).

Taken together, these findings demonstrate that multivalent TUmRNA vaccines can enhance BCG-mediated protection against pulmonary *M. tuberculosis* infection in mice and are associated with increased antigen-specific cytokine responses at the primary site of infection.

### TUmRNA_G_ protects guinea pigs against *M. tuberculosis* infection

Having demonstrated the immunogenicity, tolerability and efficacy of our multivalent mRNA vaccine candidates in mice, we next evaluated their performance in guinea pigs, a model that more closely recapitulates key features of human tuberculosis pathology^54,55^. Groups of eight outbred Dunkin-Hartley guinea pigs were immunized three times at three-week intervals with 12 µg of TUmRNA_6_, 18 µg of TUmRNA_9_, or PBS, or received a single BCG immunization on day zero (**Figure 7A**). The vaccines were well tolerated, with no apparent health concerns observed. Following aerosol *M. tuberculosis* challenge, disease progression was more pronounced in PBS-treated and TUmRNA_6_-vaccinated animals, whereas TUmRNA_9_- and BCG-vaccinated animals maintained better health overall (**Figure S15**). Early mortality was observed between days 24 and 28 post-infection in two animals from both the PBS control and TUmRNA_6_ groups. One animal in the TUmRNA_9_ group required early euthanasia at day 27 post-infection. The remaining animals were euthanized on day 28 post-infection for assessment of bacterial burden and pulmonary lesions (**Figure 7A**).

**Figure 7.**
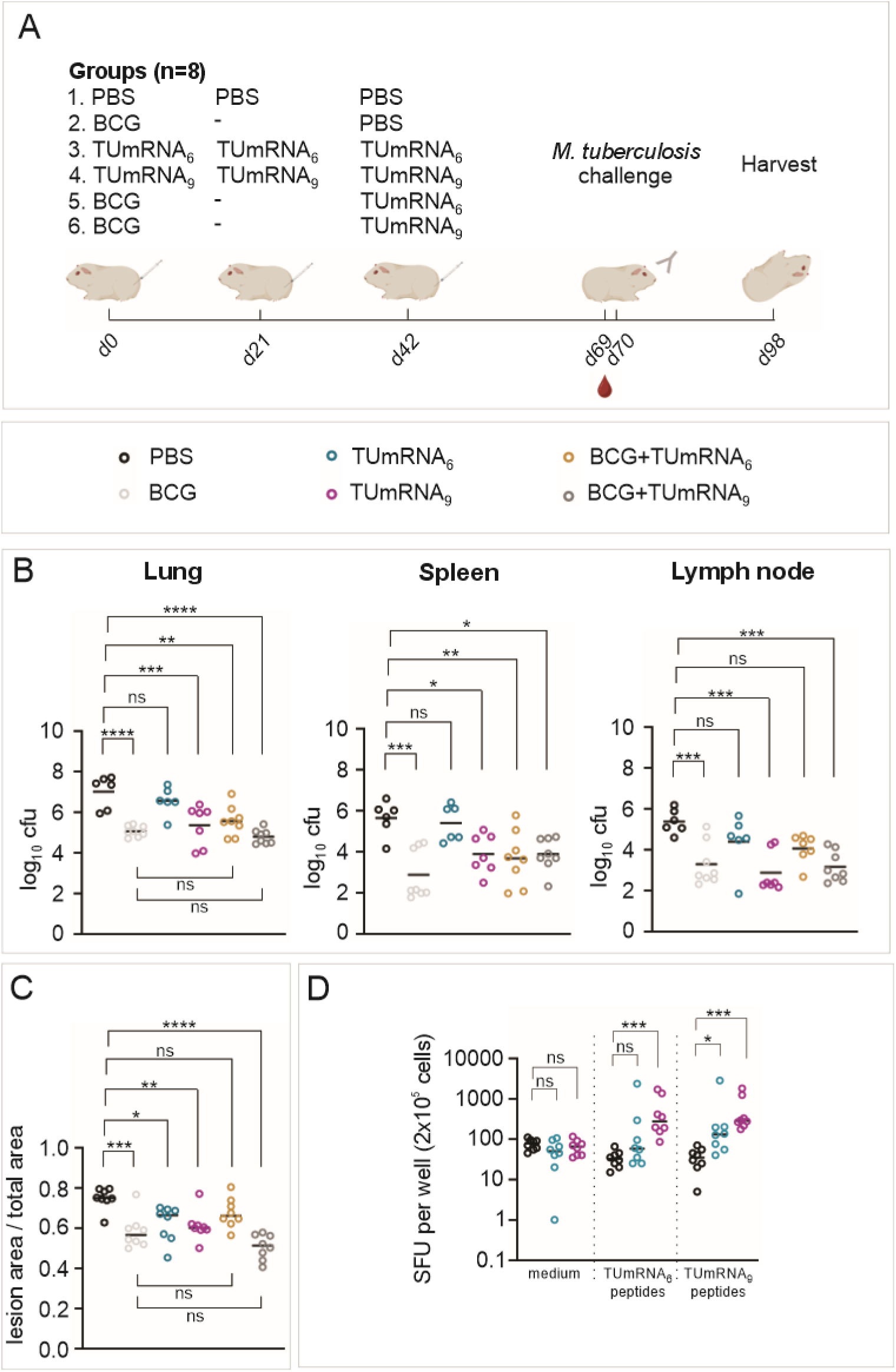
TUmRNA_G_ protects guinea pigs from *M. tuberculosis* infection. **A)** Immunization, challenge, and sample collection schedule used to evaluate the protective effects of TUmRNA_6_ and TUmRNA_9_ administered alone or following BCG priming. The schematic was generated using BioRender and CorelDRAW Graphics Suite 2026. **B)** *M. tuberculosis* burden in the lungs, spleens and mediastinal lymph nodes of vaccinated guinea pigs following aerosol challenge. Circles represent the log_10_ bacterial counts in individual guinea pigs and lines indicate means (n=6-8). **C)** Pulmonary lesion burden in vaccinated guinea pigs following *M. tuberculosis* challenge. Lesion burden was quantified as the percentage of total lung area occupied by tuberculous lesions (mean of three lung sections analyzed per animal). Lines indicated medians (n=8). In **B** and **C**, guinea pigs that died or were euthanized between days 24 and 27 post-infection were excluded from the analysis, including two PBS-treated, two TUmRNA_6_-vaccinated, and one TUmRNA_9_-vaccinated guinea pig. Statistical significance was determined using one-way ANOVA followed by Šidák’s multiple-comparisons test. *=p<0.05, **=<0.01, ***=p<0.001, ****=p<0.0001, ns=not significant. **D)** Antigen specific IFNγ-producing cells in PBMCs from vaccinated guinea pigs prior to challenge. Cells were restimulated with either vaccine-specific peptide pools or medium alone and analyzed by ELISpot. Circles represent mean spot-forming units (SFU) per 2×10^5^ cells from duplicate wells and lines indicate medians (n=8). Statistical significance was determined using Kruskal-Wallis test with Dunn’s multiple comparisons test. *<0.05, **=p<0.01, ***=p<0.001, ****<0.0001, ns=not significant.

TUmRNA_9_ vaccination significantly reduced bacterial burdens in lung, spleen and lymph nodes compared with PBS-treated animals, with mean reductions of 1.7 (p=0.0003), 1.8 (p=0.02), and 2.5 log_10_ CFU (p=0.0001), respectively (**Figure 7B**). The corresponding reductions in BCG-vaccinated animals were 2.0, 2.8, and 2.1 log_10_ CFU in the lungs (p<0.0001), spleen (p<0.0001), and lymph nodes (p=0.0008), respectively. In this model, TUmRNA_6_ conferred only modest protection with mean reductions of 0.5, 0.2 and 1.0 log_10_ CFU in the lungs, spleen and lymph nodes, respectively. However, both mRNA vaccines and BCG significantly reduced lung lesion burden compared with PBS-treatment, with reductions of 17% for TUmRNA_6_ (p=0.01), 18% for TUmRNA_9_ (p=0.005) and 22% for BCG (p=0.0004) (**Figure 7C**, representative images of lung sections in **Figure S15**). Taken together, these findings demonstrate that TUmRNA_9_ confers greater protection than TUmRNA_6_ in the guinea pig model of tuberculosis, reducing bacterial burden in all three organs to levels broadly comparable to those achieved with BCG.

In the same experiment, we also investigated whether TUmRNA_6_ or TUmRNA_9_ could enhance BCG-induced protection. Guinea pigs were primed with BCG and boosted with either mRNA vaccine six weeks later. In this setting, boosting with TUmRNA_6_ did not provide additional protection to BCG alone (**Figures 7B-C**). In contrast, animals receiving a TUmRNA_9_ boost exhibited the lowest pulmonary bacterial and lesion burdens of all groups. Compared with PBS-treated controls, the BCG-TUmRNA_9_ regimen reduced lung bacterial burden by a mean of 2.2 log_10_ CFU (p<0.0001) and lesion burden by 32% (p<0.0001). Relative to BCG alone, BCG-TUmRNA_9_ achieved an additional mean reduction of 0.3 log_10_ CFU (not significant) and 13% reduction in lesion burden (not significant). Although these differences relative to BCG alone did not reach statistical significance, TUmRNA_9_ boosting consistently yielded lowest bacterial and lesion burden among the vaccinated groups.

To compare vaccine-induced cellular responses elicited by TUmRNA_6_ and TUmRNA_9_ vaccination in guinea pigs, we analyzed vaccine specific IFNy responses one day prior to *M. tuberculosis* challenge. In peripheral blood mononuclear cells (PBMCs) restimulated with TUmRNA_6_- or TUmRNA_9_-specific peptide pools, cells derived from TUmRNA_9_-vaccinated guinea pigs produced significantly higher numbers of IFNγ-secreting cells than those from PBS-treated controls (**Figure 7D**). PBMCs from TUmRNA_6_-vaccinated guinea pigs showed similar increases following stimulations with both peptide pools; however, compared with PBS-treated animals, these differences reached statistical significance only after stimulation with the TUmRNA_9_-specific peptide pool.

These results indicate that both multivalent mRNA vaccines are immunogenic in guinea pigs, but TUmRNA_9_ provides greater protection against *M. tuberculosis* challenge in this model. Taken together, our findings demonstrate that antigens identified in zebrafish can be translated into protective multivalent mRNA vaccines, providing protection against tuberculosis across multiple animal models, including zebrafish, mice, and guinea pigs.

## Discussion

In this study, we combined antigen discovery in zebrafish with mRNA vaccine technology to develop multivalent tuberculosis vaccine candidates that were immunogenic and protective in both mice and guinea pigs. These findings add to the current evidence supporting mRNA vaccines as a promising platform for tuberculosis prevention^56^.

Several studies have reported encouraging immunogenicity and efficacy data for mRNA-LNP-based tuberculosis vaccines in murine models^21–24,26^, and one candidate has advanced to clinical evaluation^28^. However, antigen selection for tuberculosis vaccines remains a major challenge, largely due to the complex and dynamic nature of *M. tuberculosis* infection and the stage-dependent expression of antigens during disease progression^57,58^. While many vaccine candidates have been based on a limited number of highly immunogenic antigens, such as members of the Ag85 family, ESAT-6, and CFP-10^57,59–62^, we selected multiple antigens expressed during different stages of *M. tuberculosis* infection. Our approach differed from conventional antigen discovery strategies by using a zebrafish model to identify mycobacterial antigens that protect against primary infection and/or the reactivation of latent disease^35,37,39,40^. Based on these findings, we generated three multivalent mRNA vaccines. The protective efficacy observed in mammalian models supports the translational potential of this discovery platform and highlights the value of alternative infection models for identifying novel vaccine targets.

mRNA vaccines induce both cellular and humoral immune responses^63,64^. Previous studies on mRNA-based tuberculosis vaccines have reported increased frequencies of antigen-specific IFNγ-, TNF-, and IL2-producing CD4+ and CD8+ T cells, as well as polyfunctional T cells following vaccination^22–26,28^. These cells have also frequently been associated with protection against *M. tuberculosis* in other mouse studies^65–68^. In our study, TUmRNA vaccines were immunogenic in mice and guinea pigs and induced antigen-specific IFNγ responses. More detailed characterization in mice showed that TUmRNA_6_ elicited a broad cytokine profile indicating a general immune system activation without evident Th1/Th2 skew. We also saw an expansion of central and effector memory T cell populations related to tissue and long-term memory formation^69^. Importantly, simultaneous responses to multiple vaccine antigens were detected, indicating that the inclusion of several targets within a single vaccine did not result in obvious immunological interference. Following *in vivo* challenge, TUmRNA vaccination promoted the expansion of lung memory T cells, although the response varied between vaccine formulations. TUmRNA_6_-mediated protection was associated with increased frequencies of central memory CD8+ T cells, whereas TUmRNA_9_ vaccination preferentially expanded central memory CD4+ T cells. In BCG-primed mice, TUmRNA vaccination further enhanced antigen-specific IFNγ, TNF, and IL2 production in the lungs compared with BCG alone, while TUmRNA_6_ boosting also increased IL17A responses, a feature previously associated with improved protection against *M. tuberculosis*^70,71^. Although antigen-specific antibody responses were not systematically investigated, the detection of anti-GFP antibodies in TUmRNA_GFP_-vaccinated mice suggests that our platform is capable of inducing humoral immunity. This may be advantageous given the increasing recognition of antibodies as contributors to host defense against *M. tuberculosis*^72^.

Consistent with previous studies demonstrating the protective efficacy of mRNA-based tuberculosis vaccines in murine studies^21–24,26^, vaccination with multivalent TUmRNA formulations conferred substantial protection against acute *M. tuberculosis* infection in mice. TUmRNA_4_ and TUmRNA_6_ reduced pulmonary bacterial burden and lung pathology, reaching levels of protection comparable to those achieved with BCG. Although TUmRNA_9_ also reduced bacterial burden in the lungs (with statistical significance achieved only when the two-dose and three-dose regimens were combined), the inclusion of three additional antigens did not further enhance protection relative to TUmRNA_6_ in mice. Notably, TUmRNA vaccination also reduced bacterial burden in the spleen, and in guinea pigs, also in the mediastinal lymph nodes, indicating an ability to limit bacterial dissemination beyond the primary site of infection. This is particularly relevant for young children exposed to *M. tuberculosis*, in whom the disease can progress rapidly to severe, disseminated forms involving extrapulmonary tissues^73–75^. Our findings suggest that these multivalent mRNA vaccine candidates may have value in limiting extrapulmonary spread of infection in this age group while avoiding the safety concerns associated with live BCG vaccination.

TUmRNA_6_ enhanced BCG-mediated protection in mice when administered as a booster. Given that BCG remains the only licensed tuberculosis vaccine and is routinely administered worldwide^76^, mRNA-based booster strategies offer an attractive opportunity to build on existing BCG-induced immunity. This is particularly relevant given the large global population with latent *M. tuberculosis* infection, including individuals previously vaccinated with BCG^1,77^. Importantly, several antigens included in our vaccine candidates were specifically selected based on their preferential expression during reactivation and their ability to prevent mycobacterial reactivation in zebrafish. This rationale, together with the observed enhancement of BCG-mediated protection, supports further investigation of TUmRNA vaccines as boosters designed to strengthen existing immunity and potentially prevent or reduce reactivation of latent tuberculosis.

To date, evaluation of mRNA-based tuberculosis vaccines has been largely restricted to murine models, despite the long-established value of guinea pigs in preclinical tuberculosis vaccine testing^54,55^. To our knowledge, this study provides the first demonstration of protective efficacy of an mRNA tuberculosis vaccine in the guinea pig model. TUmRNA_9_ reduced bacterial burden in multiple organs, with effects comparable to those of BCG vaccination, and reduced pulmonary lesion severity, demonstrating substantial protection against disease. Notably, vaccine performance differed between species: whereas TUmRNA_6_ showed great protective efficacy in mice, TUmRNA_9_ was more protective in guinea pigs. Although the mechanisms underlying these differences were not investigated in this study, they may reflect species- or strain-specific variation in immune responses, disease progression, and antigen recognition, underscoring the importance of evaluating vaccine candidates across multiple preclinical models.

Unlike live attenuated vaccines, mRNA vaccines are non-replicating and contain no infectious agent capable of reverting to a pathogenic form, providing a favorable safety profile, particularly in immunocompromised populations^78^. The clinical experience gained from billions of administered mRNA vaccine doses has further demonstrated the safety and tolerability of this platform across diverse populations^20,78,79^, including children^80,81^. In the present study, the multivalent TUmRNA vaccine candidates were well tolerated in all three animal species tested.

In conclusion, we demonstrate that multivalent mRNA vaccines encoding antigens identified in zebrafish model induce robust immune responses and confer protection against mycobacterial infection in three animal species, with efficacy comparable to that of BCG. The ability of TUmRNA_6_ to match BCG efficacy and enhance BCG-mediated protection, together with the strong performance of TUmRNA_9_ in guinea pigs, highlights the promise of this approach. Based on the ability of TUmRNA vaccines to protect against *M. marinum* infection in zebrafish, multivalent mRNA vaccines may also offer protection against non-tuberculous mycobacteria. Altogether, these findings support the continued development of multivalent mRNA tuberculosis vaccines and provide a strong rationale for further evaluating TUmRNA vaccine candidates.

### Limitations of the study

Multiple animal models, including mice, other rodent species, and non-human primates, have been used to evaluate tuberculosis vaccine efficacy; however, none fully recapitulates human tuberculosis or reliably predicts vaccine performance in humans. In this study, we employed two widely used preclinical models, a low-dose aerosol challenge in C57BL/6 mice, and the guinea pig model, to assess the immunogenicity and protective efficacy of multivalent TUmRNA vaccine candidates. Future studies incorporating additional mouse genetic backgrounds, diverse *M. tuberculosis* strains, and alternative challenge systems, such as the ultra-low-dose aerosol model, which more closely mimics natural human exposure^82^, would provide a more comprehensive assessment of vaccine efficacy. Furthermore, immune responses and protection were only evaluated at a single experimental time point, limiting our ability to fully characterize the kinetics, durability, and mechanistic basis of vaccine-induced immunity.

Reactivation of latent infection accounts for a substantial proportion of active tuberculosis cases worldwide and represents a critical unmet need in vaccine development^1,77^. By incorporating antigens shown to be protective against reactivation of latent mycobacterial infection in zebrafish into the TUmRNA_6_ and TUmRNA_9_ vaccine formulations, we sought to address this challenge. However, due to the lack of an animal model that fully recapitulates human latency and reactivation, the capacity of these vaccines to protect against reactivation remains to be elucidated.

## Resource availability

Requests for further information and resources should be directed to, and will be fulfilled by, the lead contact, Prof. Mika Rämet

## Acknowledgements

We acknowledge the Biocenter Finland (BF) and Tampere Zebrafish Core Facility, Tampere Genomics Facility, Tampere University Flow Cytometry Facility (TFCF), and Tampere preclinical Facility for their service. We thank the Veterinary Pathology Diagnostic service at the University of Sydney for histological sample processing and evaluation. We also thank Tanja Salomaa and Niina Ahvenainen for technical assistance. We also acknowledge all the funding sources including the Jane and Aatos Erkko Foundation (M.R., B.M.S.), the Sigrid Jusélius Foundation (M.R.), the Tampere Tuberculosis Foundation (M.R., A.K.S., T.A.A.I., I.S.J., S.V.), the Orion Research Foundation (A.K.S., T.A.A.I), the Väinö and Laina Kivi Foundation (A.K.S., T.A.A.I), Foundation of the Finnish Anti-Tuberculosis Association (A.S., T.A.A.I), Finnish Society for Study of Infectious Diseases (T.A.A.I), the Ida Montin Foundation (T.A.A.I.), the Alma and K.A. Snellman Foundation (M.J.), Competitive State Research Financing of the Expert Responsibility Area of Fimlab Laboratories (grant: X51409) (I.S.J), Research Council of Finland (grant: 3121369146) (I.S.J), the Competitive State Research Financing of the Expert Responsibility Area of NordLab (grant: KT0016) (I.S.J).

## Author contributions

Conceptualization: MR, BMS, AS, ISJ, BKP, DFA

Data Curation: AS, TAAI, MTN, BMS, DFA, LH, ORM, AE

Formal Analysis: AS, TAAI, MTN, LK, MJ, SV, BMS, DN, RJKE, DFA, LH, ORM, AE, ISJ, MR

Funding acquisition: MR, BMS

Investigation: AS, TAAI, MTN, LK, LM, DN, DFA, LH, ORM, AE, RJKE, MG-R, MJ, SV, BKP, BMS

Methodology: AS, MTN, LK, DFA, LH, ORM, AE, BKP, BMS, ISJ, MR

Validation: AS, BMS

Visualization: AS, TAAI, DFA, ORM

Project administration: MR, BMS, BKP, AS

Supervision: MR, BMS, BKP, AS

Writing – original draft: AS, TAAI

Writing – review C editing: AS, TAAI, MTN, LK, LM, DN, DFA, LH, ORM, AE, RJKE, MG-R, MJ, SV, ISJ, BKP, BMS, MR

## Declaration of generative AI and AI-assisted technologies in the manuscript preparation process

During the preparation of this work, the authors used Microsoft 365 Copilot or ChatGPT for proof-reading the text. After using this tool, the authors reviewed and edited the content as needed and take full responsibility for the content of the published article.

## Supplemental Information

Document S1. Supplementary Figures S1-S15

Document S2. Supplementary Tables S1-S4

