## Supplementary Figures for "Multivalent mRNA tuberculosis vaccines protect against mycobacterial infection in zebrafish, mice and guinea pigs"

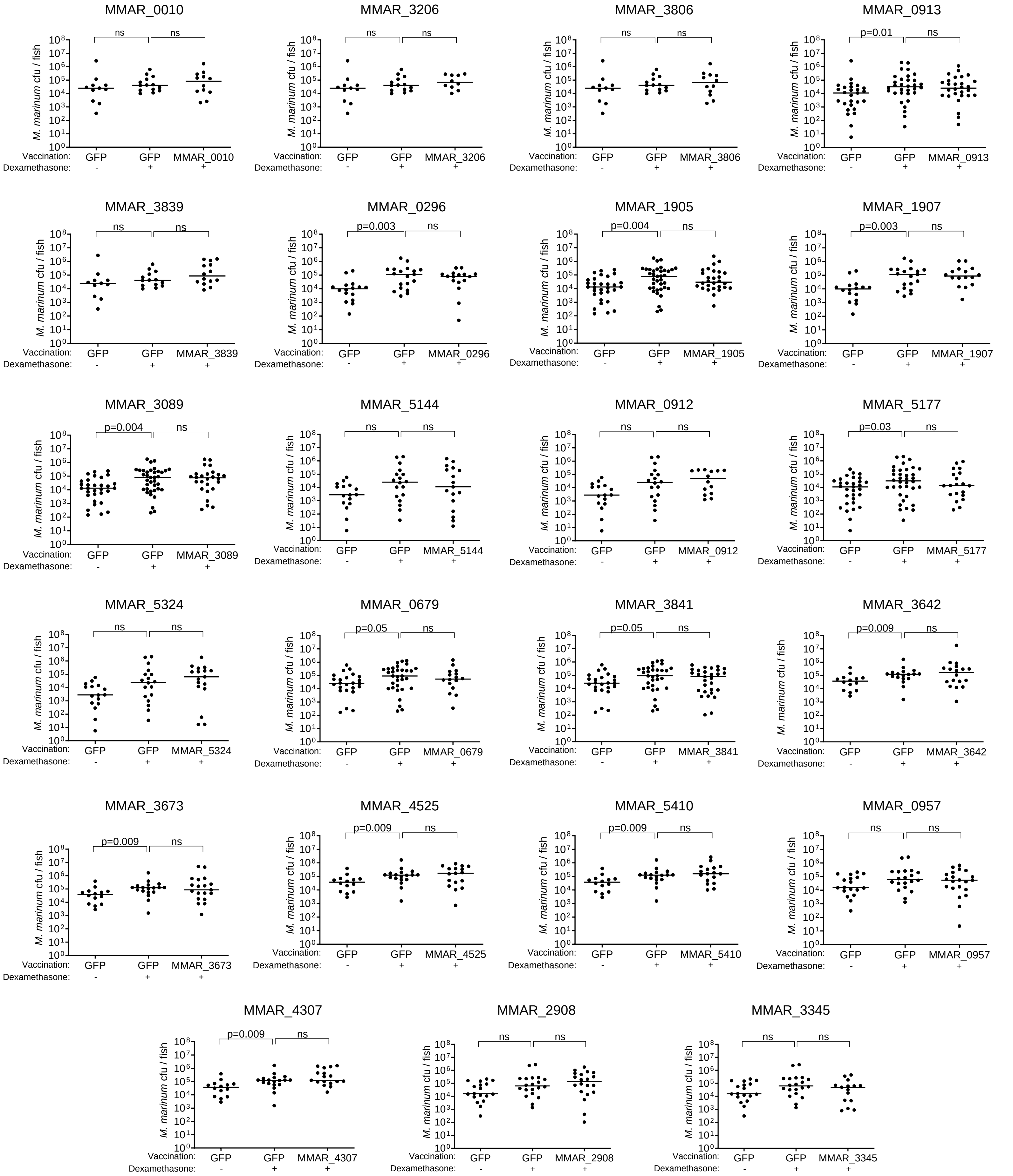


**Figure S1.** Effect of 23 screened DNA vaccine candidates on the reactivation of latent *M. marinum* infection in zebrafish. Latent *M. marinum* infection was established in zebrafish by low-dose intraperitoneal injection. Five weeks post-infection, fish were vaccinated, and 5 weeks later, reactivation was induced with dexamethasone. Bacterial burden was measured after 4 weeks of treatment. The figure shows the bacterial counts in zebrafish vaccinated with DNA vaccines encoding individual *M. marinum* genes or with the control vaccine encoding GFP. Data represent results from one or two independent experiments (n= 11-36 per group). Dots indicate colony forming units (CFU) in single fish and lines indicate medians. Statistical significance was calculated using two-tailed Mann-Whitney test. ns=not significant.

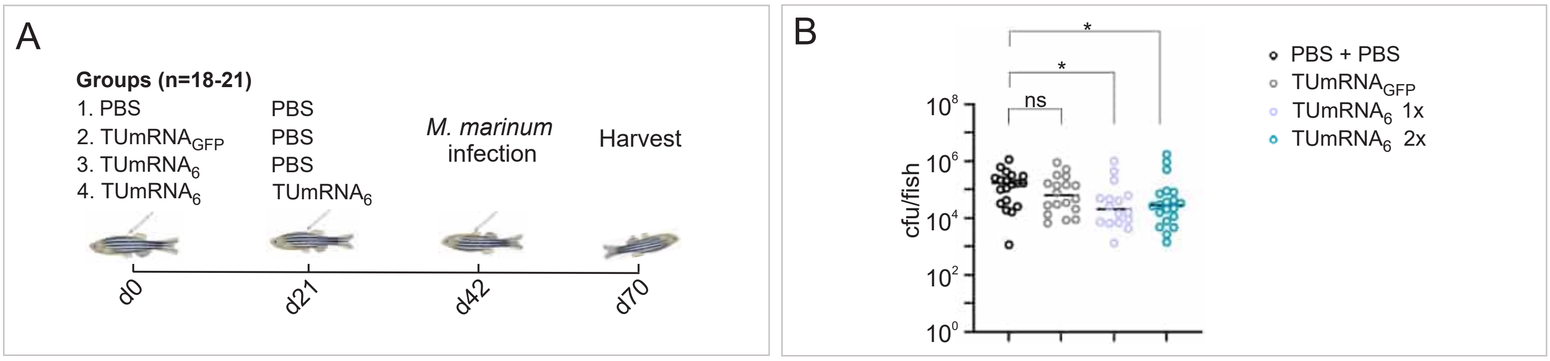


**Figure S2. An mRNA vaccine encoding six *M. tuberculosis* antigens protects zebrafish from *M. marinum* infection. A)** Immunization and challenge schedule of the study evaluating the protective effect of $TUmRNA_6$ against *M. marinum* infection in zebrafish. The schematic was generated using BioRender and CorelDRAW Graphics Suite 2026. **B)** Vaccination with $TUmRNA_6$ reduces *M. marinum* burden in adult zebrafish. Circles represent bacterial counts in individual fish alive at the time of sampling, lines indicate medians (n=18-21). Fish that were euthanized between days 42 and 70 were excluded from the analysis, but included one fish in PBS group, two fish in $TUmRNA_6$ 1x group, and one fish in $TUmRNA_6$ 2x group. Statistical significance was calculated using Mann-Whitney test. *=p<0.05, ns=not significant.

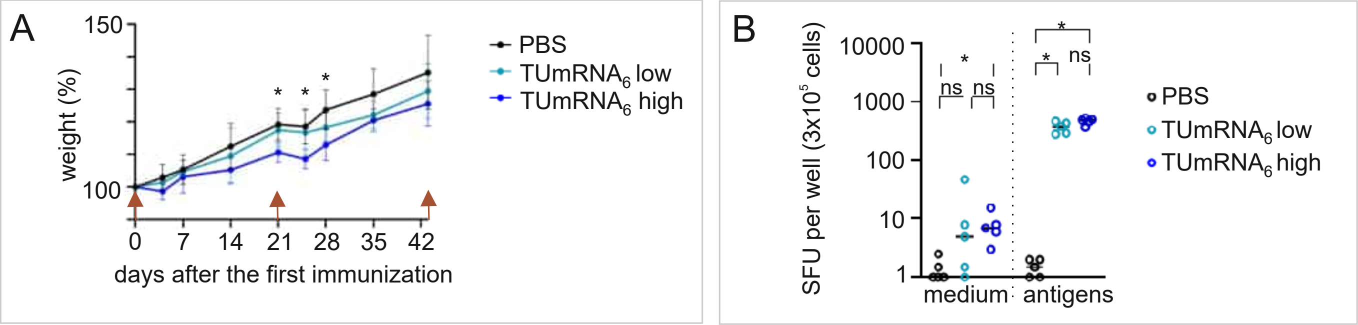


**Figure S3. High and low doses of TUmRNA$_6$ induce comparable antigen-specific IFNγ responses in mice.** Mice were immunized three times at three-week intervals with a high dose (12 µg) or a low dose (2 µg) of TUmRNA$_6$ or PBS. **A)** Body weight of mice immunized with PBS or TUmRNA$_6$ expressed as percentage of baseline weight (mean + SD, n = 5); red arrows indicate vaccination time points. Statistical significance was calculated using the Mann–Whitney test, and the asterisks depict significant differences (*=p<0.05) between PBS and TUmRNA$_6$ high dose groups at indicated time points. **B)** Splenocytes were harvested three days post final immunization, restimulated for 16 h with TUmRNA$_6$-specific peptide pool or medium and analyzed for IFNγ response by ELISpot. Circles represent mean spot-forming units (SFU) per 3x10$^5$ cells from duplicate wells and lines indicate medians (n=5). Statistical significance was calculated using the Mann-Whitney test with Bonferroni-Holm correction for multiple comparisons. *=p-value below the threshold determined by Bonferroni-Holm correction, ns=not significant.

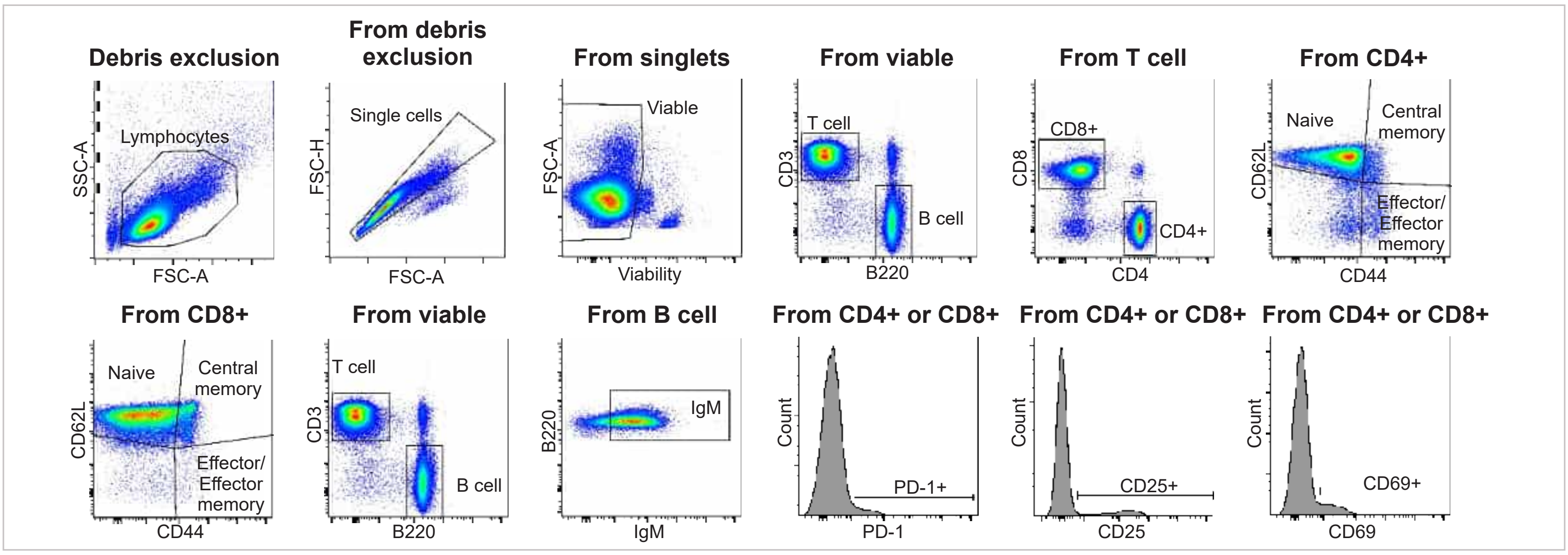


**Figure S4. Gating strategy in flow cytometry analysis.** Representative gating strategy for the selected populations in the inguinal lymph node.

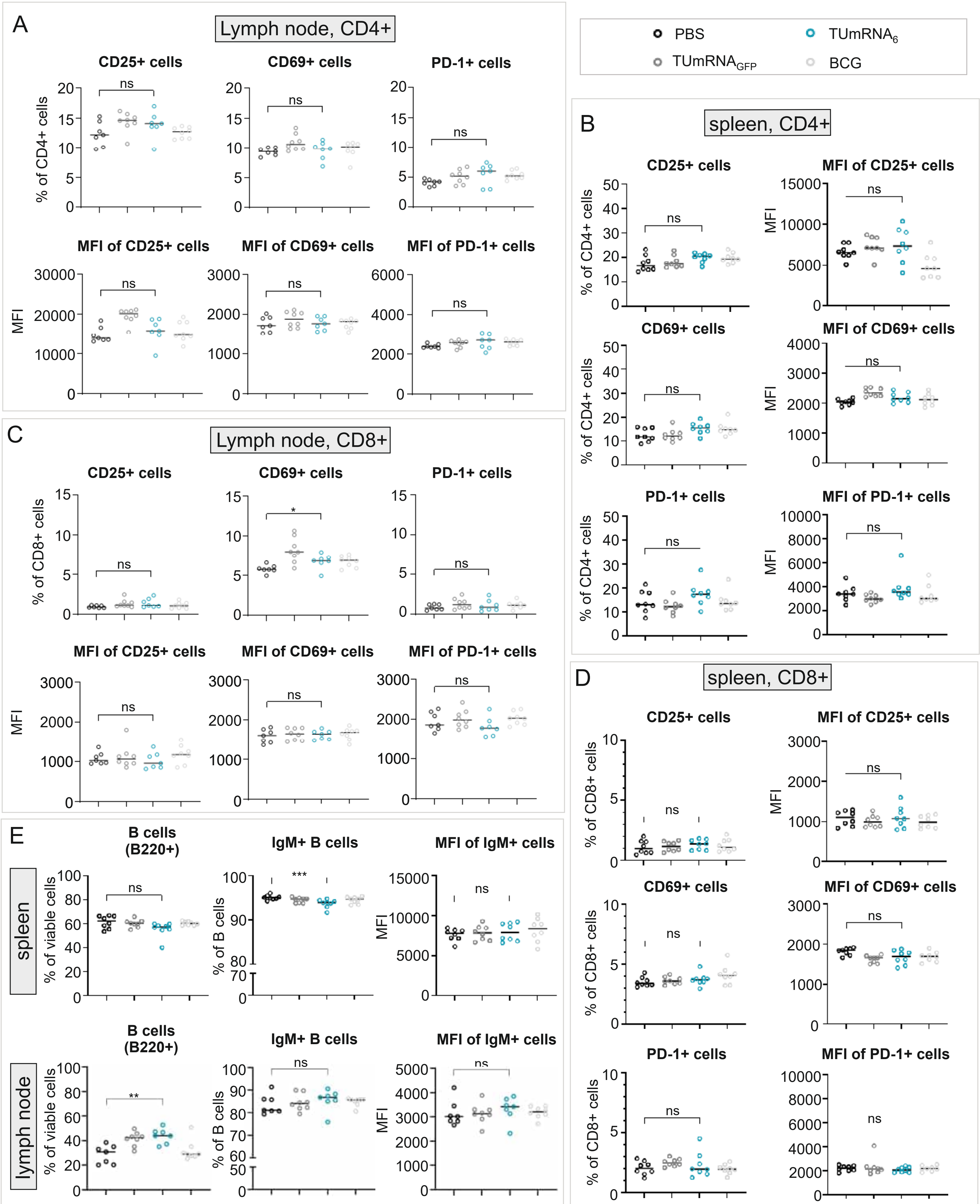


**Figure S5. Vaccination with $TUmRNA_6$ has little effect on the prevalence of B cells and the activation of T cells in mice.** Spleen and inguinal lymph nodes were collected three days post final vaccination and selected lymphocyte populations were analyzed by flow cytometry. The frequencies of CD4+ cells expressing CD25+, CD69+ and PD-1+ in (**A**) lymph nodes and (**B**) spleen, and the expression levels of the markers as median fluorescence intensity (MFI). The frequencies of CD8+ cells expressing CD25+, CD69+ and PD-1+ in (**C**) lymph nodes and (**D**) spleen, and the expression levels of the markers as median fluorescence intensity (MFI). **E**) Frequencies of total B cells (B220+) and IgM+ B cells in the spleen and lymph nodes, and the IgM expression level on IgM+ cells as median fluorescence intensity (MFI). Circles represent individual mice and lines depict medians (n=7-8). Statistical significance was calculated using Mann-Whitney test. *=p<0.05, **=p>0.01, ***p<0.001, ns=not significant.

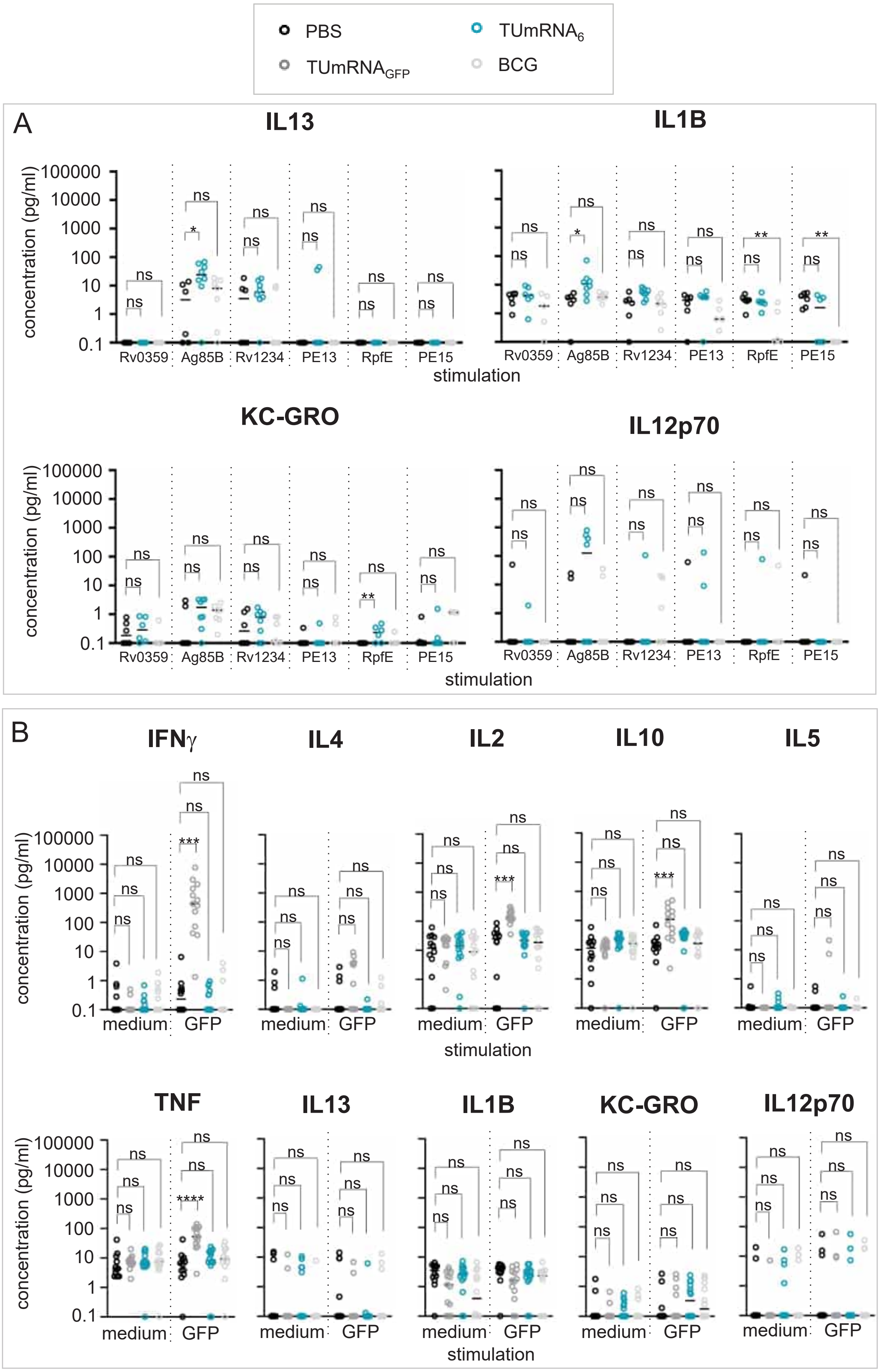


**Figure S6. Splenocytes from TUmRNA$_6$ vaccinated mice produce diverse cytokines following antigen restimulation.** Splenocytes were isolated three days after the final immunization and stimulated with antigen-specific peptide pools (2 µg/ml) or medium for 68 h. After stimulation, concentrations of multiple Th1/Th2 cytokines in culture supernatants were quantified using a multiplex cytokine assay. **A)** Concentrations of IL13, IL1B, KC-GRO and IL12p70 after stimulation with indicated peptide pools. **B)** Concentrations of IFNγ, IL4, IL2, IL10, IL5, TNF, IL13, IL1B, KC-GRO, and IL12p70 after stimulation with GFP peptide library or medium. In **A** and **B**, circles represent individual measurements, and lines indicate medians (n=5-8, one BCG-vaccinated mouse was excluded for health issues unrelated to vaccination). Statistically significant differences were calculated using the Kruskal-Wallis test with Dunn's multiple-comparisons test. *=p<0.05, **=p<0.01, ***p<0.001, ns=not significant.

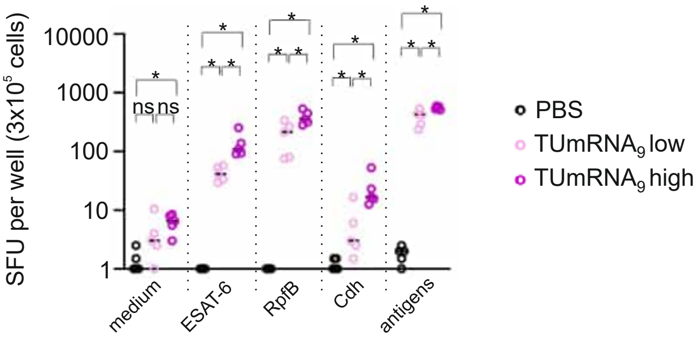


**Figure S7. ESAT-6, RpfB, and Cdh are immunogenic in mice inducing antigen-specific IFNγ response by splenocytes.** Mice were immunized three times at three-week intervals with either high (18 µg) or a low (2 µg) of $TUmRNA_9$ or PBS. Splenocytes were harvested three days post final immunization, restimulated for 16 h with indicated peptide pools or medium and analyzed for IFNγ response by ELISpot. Circles represent mean spot-forming units (SFU) per $3x10^5$ cells from duplicate wells and lines indicate medians (n=5). Statistical significance was calculated using the Mann-Whitney test with Bonferroni-Holm correction for multiple comparisons. *=p-value below the threshold determined by Bonferroni-Holm correction, ns=not significant.

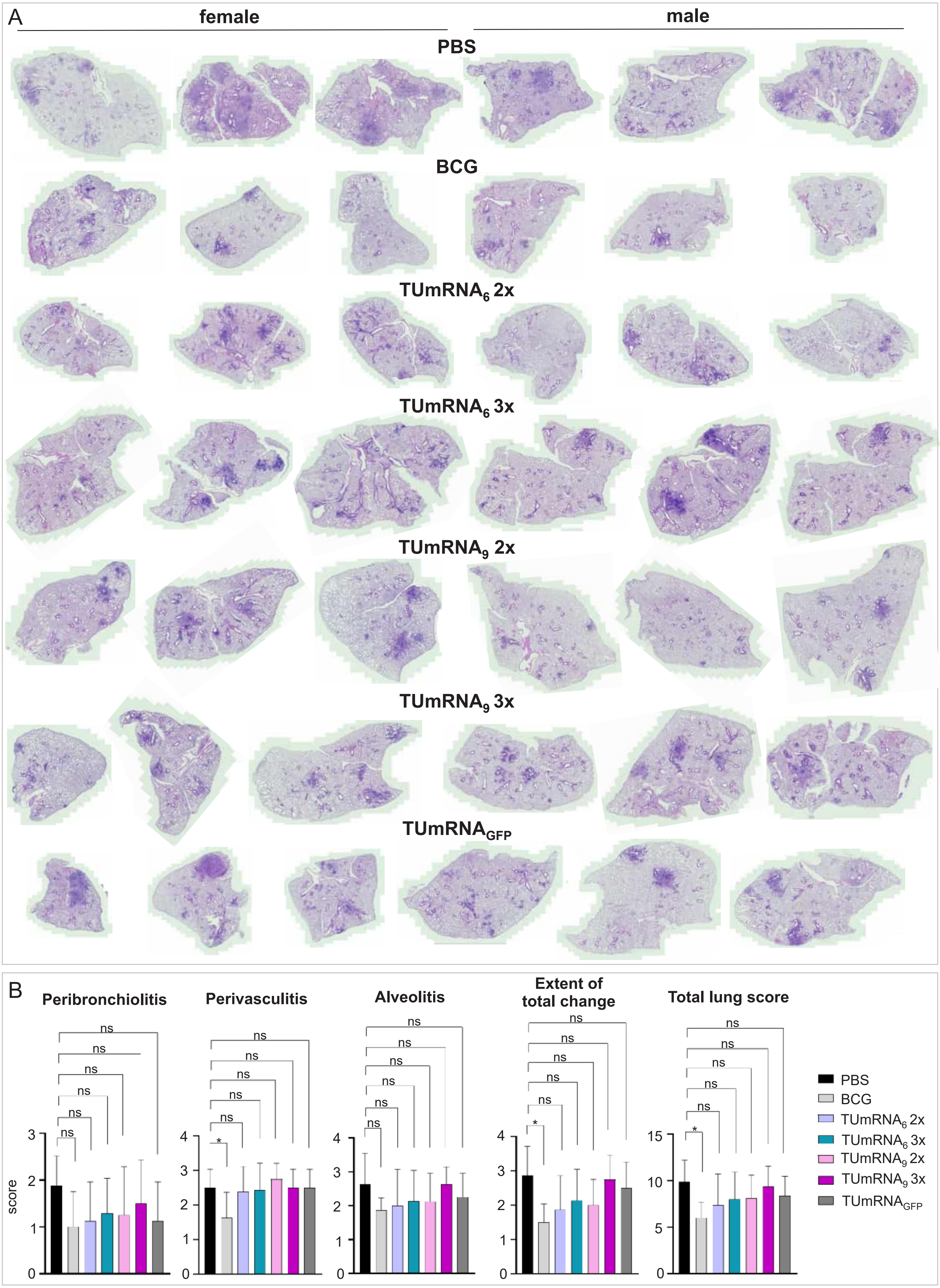
A
female
male
PBS
BCG
TUmRNA6 2x
TUmRNA6 3x
TUmRNA9 2x
TUmRNA9 3x
TUmRNAGFP
B
Peribronchiolitis
Perivasculitis
Alveolitis
Extent of total change
Total lung score
score
ns
*
PBS
BCG
TUmRNA6 2x
TUmRNA6 3x
TUmRNA9 2x
TUmRNA9 3x
TUmRNAGFP

**Figure S8. Mouse lung histopathology following two or three doses of $TUmRNA_6$ or $TUmRNA_9$ and *M. tuberculosis* challenge.** Mice were immunized two or three times with $TUmRNA_6$ or $TUmRNA_9$ at three-week intervals and challenged with *M. tuberculosis* four weeks after final immunization. Five weeks post-challenge, the middle right lung lobe was collected, sectioned, and stained with H&E. **A)** Representative lung sections from three females and three males. **B)** Lung sections were scored for peribronchiolitis, perivasculitis, alveolitis, and overall pathological changes. Individual scores were summed to generate a total lung pathology score. Data are presented as mean ± standard deviation (n=8). Statistical significance was calculated using one-way ANOVA followed by Tukey's multiple-comparison test. *<0.05, ns=not significant.

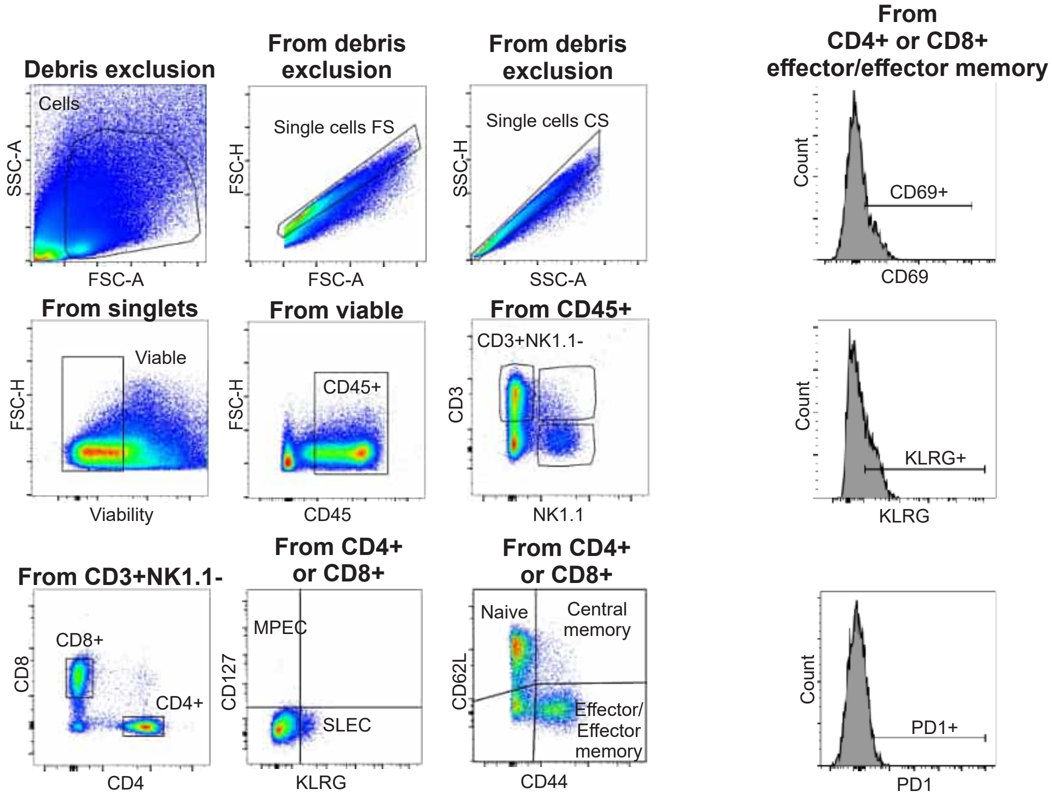


**Figure S9. Gating strategy in flow cytometry analysis.** Representative flow cytometry gating strategy for the selected populations in the lung.

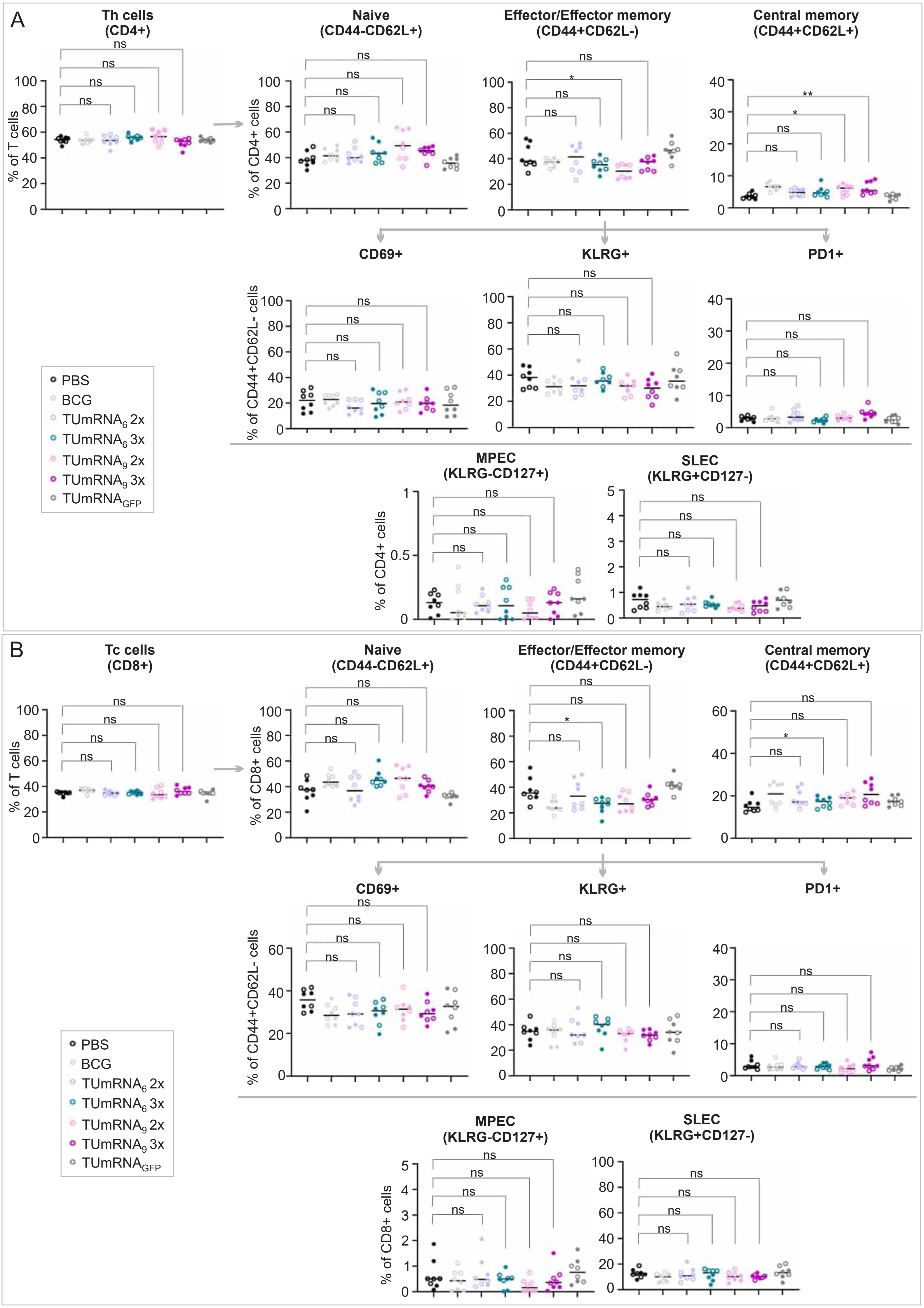

A
Th cells (CD4+)
% of T cells
Naive (CD44-CD62L+)
% of CD4+ cells
Effector/Effector memory (CD44+CD62L-)
Central memory (CD44+CD62L+)
CD69+
KLRG+
PD1+
% of CD44+CD62L- cells
MPEC (KLRG-CD127+)
SLEC (KLRG+CD127-)
% of CD4+ cells
ns
*
**
PBS
BCG
TUmRNA$_6$ 2x
TUmRNA$_6$ 3x
TUmRNA$_9$ 2x
TUmRNA$_9$ 3x
TUmRNA$_{GFP}$
B
Tc cells (CD8+)
% of T cells
Naive (CD44-CD62L+)
% of CD8+ cells
Effector/Effector memory (CD44+CD62L-)
Central memory (CD44+CD62L+)
CD69+
KLRG+
PD1+
% of CD44+CD62L- cells
MPEC (KLRG-CD127+)
SLEC (KLRG+CD127-)
% of CD8+ cells
PBS
BCG
TUmRNA$_6$ 2x
TUmRNA$_6$ 3x
TUmRNA$_9$ 2x
TUmRNA$_9$ 3x
TUmRNA$_{GFP}$

**Figure S10. Effect of two and three doses of $TUmRNA_6$ or $TUmRNA_9$ on T cell responses to *M. tuberculosis* in mouse lung**. Mice were immunized two or three times with either $TUmRNA_6$ or $TUmRNA_9$ at three-week intervals and challenged with *M. tuberculosis* four weeks after final immunization. Five weeks post-challenge, T cells were analyzed by flow cytometry. Frequency of CD4+ (**A**) and CD8+ (**B**) T cell subsets in mouse lung. The image shows the percentages of naive (CD44-CD62L+), effector memory (CD44+CD62L-) and central memory (CD44+CD62L+) T cells, effector memory cells expressing activation markers CD69, KLRG and PD-1, memory precursor effector cells (MPEC; KLRG-CD127+), and short-lived effector cells (SLEC; KLRG+CD127-). In panels **A** and **B**, circles and dots represent female and male mice, respectively, and lines depict medians (n=8). Statistical significance was calculated using Kruskal-Wallis test with Dunn's multiple-comparisons test. *=p<0.05, **=p<0.01, ns=not significant.

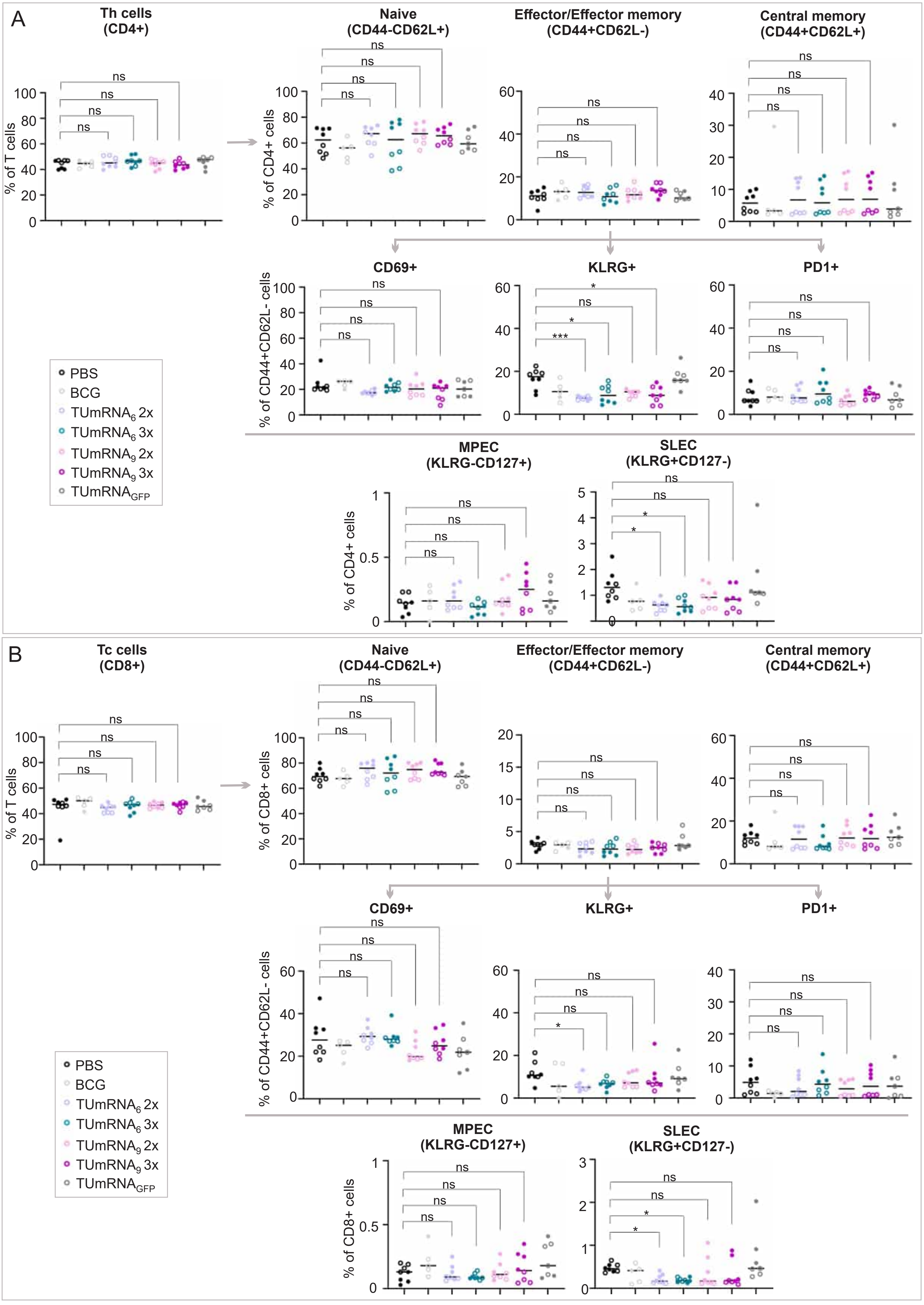

A
Th cells (CD4+)
Naive (CD44-CD62L+)
Effector/Effector memory (CD44+CD62L-)
Central memory (CD44+CD62L+)
% of T cells
% of CD4+ cells
CD69+
KLRG+
PD1+
% of CD44+CD62L- cells
MPEC (KLRG-CD127+)
SLEC (KLRG+CD127-)
ns
*
***
PBS
BCG
TUmRNA6 2x
TUmRNA6 3x
TUmRNA9 2x
TUmRNA9 3x
TUmRNAGFP
B
Tc cells (CD8+)
Naive (CD44-CD62L+)
Effector/Effector memory (CD44+CD62L-)
Central memory (CD44+CD62L+)
% of CD8+ cells
CD69+
KLRG+
PD1+
MPEC (KLRG-CD127+)
SLEC (KLRG+CD127-)

**Figure S11. Effect of two and three doses of TUmRNA$_6$ or TUmRNA$_9$ on T cell responses to *M. tuberculosis* in mouse lymph nodes.** Mice were immunized two or three times with TUmRNA$_6$ or TUmRNA$_9$ at three-week intervals and challenged with *M. tuberculosis* four weeks after final immunization. Five weeks post-challenge, T cells were analyzed by flow cytometry. Frequencies of CD4+ (**A**) and CD8+ (**B**) T cell subsets in lymph nodes. The image shows the percentages of naive (CD44-CD62L+), effector memory (CD44+CD62L-) and central memory (CD44+CD62L+) T cells, effector memory cells expressing activation markers CD69, KLRG and PD-1, memory precursor effector cells (MPEC; KLRG-CD127+) and short-lived effector cells (SLEC; KLRG+CD127-). In panels **A** and **B**, circles and dots represent female and male mice, respectively, and lines indicate medians (n=8). Statistical significance was calculated using Kruskal-Wallis test with Dunn's multiple-comparisons test. *=p<0.05, ***=p<0.001, ns=not significant.

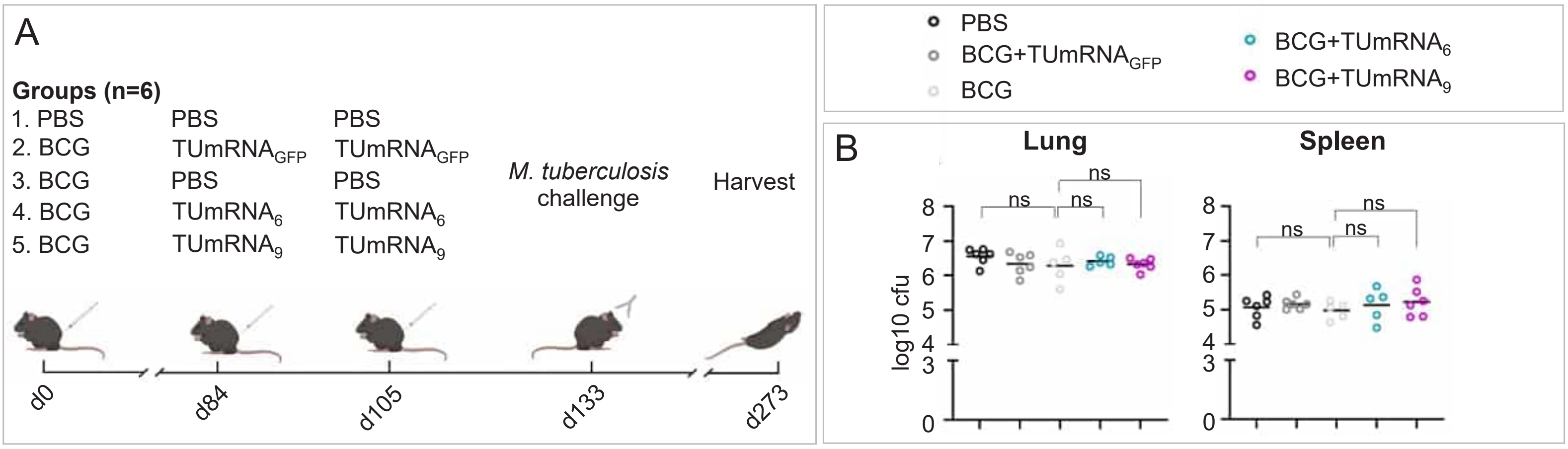


**Figure S12. Boosting BCG with TUmRNA$_{6}$ or TUmRNA$_{9}$ does not confer long-term protection against *M. tuberculosis* infection. A)** Immunization and challenge schedule of the mouse study evaluating the ability of TUmRNA$_{6}$ and TUmRNA$_{9}$ to induce longer-term protection when used as BCG boosters. The schematic was generated using BioRender and CorelDRAW Graphics Suite 2026. **B)** *M. tuberculosis* burden in lungs and spleens. Circles represent the $log_{10}$ bacterial counts in individual mice and lines indicate means (n=5-6). Statistical significance was calculated using Welch's t-test. ns= not significant.

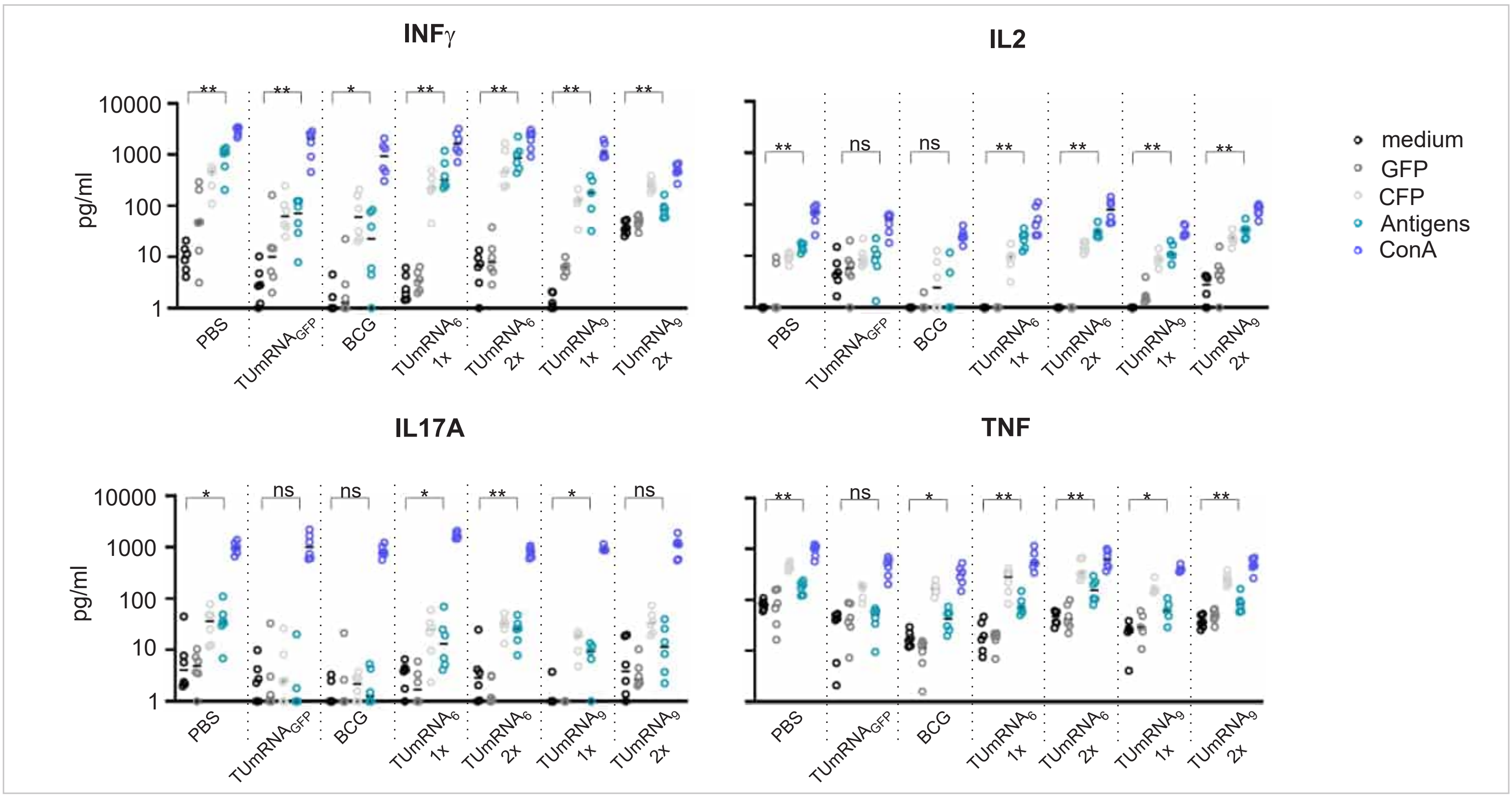


**Figure S13. TUmRNA$_6$ and TUmRNA$_9$ induce antigen-specific cytokine response in the lungs of BCG-primed mice.** BCG-primed mice were vaccinated once or twice with TUmRNA$_6$ or TUmRNA$_9$, and four weeks after final immunization, mice were challenged with *M. tuberculosis*. Four weeks later, lung cells were harvested and 1x10$^5$ cells were restimulated for 72 h with medium, a GFP peptide pool, *M. tuberculosis* culture filtrate proteins (CFP), a TUmRNA$_6$-speicific peptide pool (antigens), or Concanavalin A (ConA). IFNγ, TNF, IL2 and IL17A concentrations in culture medium were measured by cytometric bead array. Figure represents the same data as in Figure 6C. Circles indicate mean values from duplicate measurements, and lines represent group medians (n=6). Values below 1 were set to 1 for visualization purposes. Statistical significance was calculated using the Mann-Whitney test. *=p<0.05, **=p<0.01, ns=not significant.

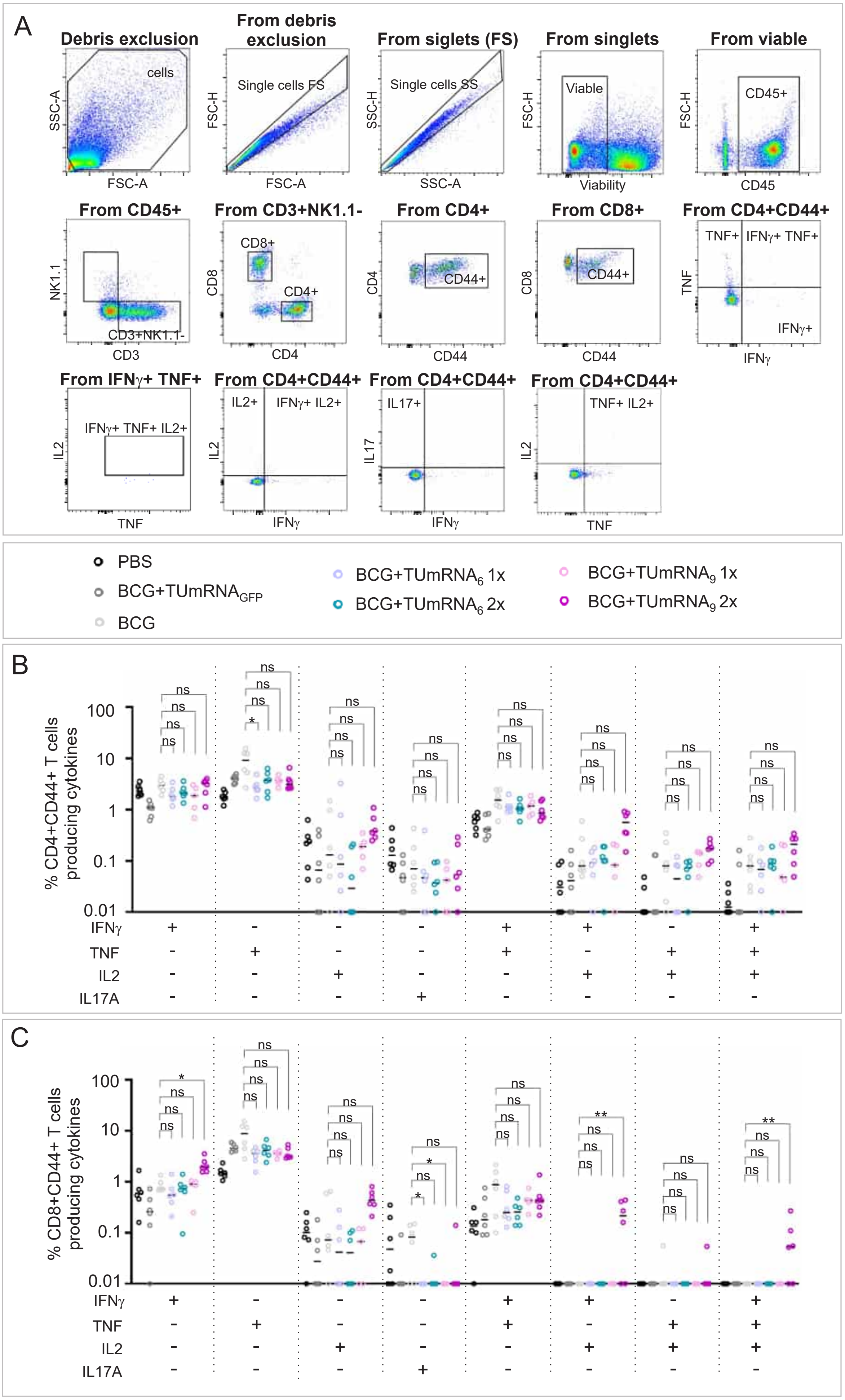


**Figure S14. Frequencies of polyfunctional cytokine-producing T cells in the lungs of BCG-primed, TUmRNA$_6$ or TUmRNA$_9$-boosted and *M. tuberculosis* challenged mice.** BCG-primed mice were vaccinated once or twice with TUmRNA$_6$ or TUmRNA$_9$, and four weeks after final immunization, mice were challenged with *M. tuberculosis*. Four weeks later, lung cells were isolated and stimulated overnight with *M. tuberculosis* culture filtrate proteins. Polyfunctional cytokine-producing T cells were quantified by intracellular cytokine staining. **A)** Representative gating strategy for selected cell populations in a mouse vaccinated with TUmRNA$_9$. Gating strategy is only shown for CD4+ Cells. **B)** Frequencies of CD4+ T cells producing indicated cytokines. **C)** Frequencies of CD8+ T cells producing indicated cytokines. In **B** and **C**, circles represent individual animals and lines indicate medians (n=5-6, one TUmRNA$_9$ 1x mouse was excluded from the study due to an injury unrelated to vaccination). Values below 0.01 were set to 0.01 for visualization purposes. Statistical significance was calculated using the Kruskal-Wallis test with Dunn's multiple-comparisons test. *=p<0.05,

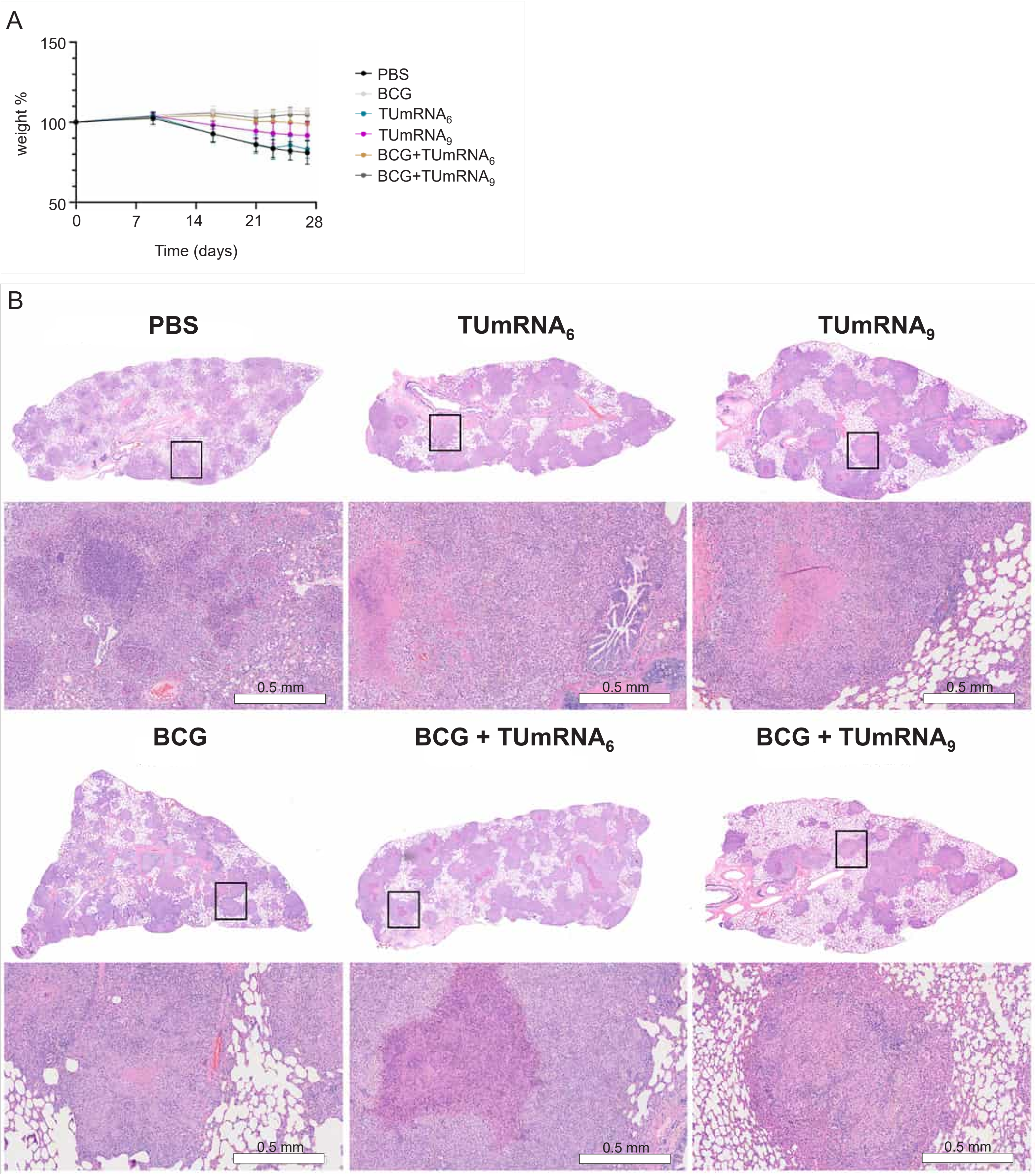


**Figure S15. Guinea pig body weights and lung histology.** Guinea pigs received PBS, $TUmRNA_6$ or $TUmRNA_9$ with or without BCG-priming, or BCG alone, and were subsequently challenged with *M. tuberculosis.* **A)** Guinea pig body weights following *M. tuberculosis* challenge. Values are expressed as a percentage of baseline body weight (mean + SD, n=8). Two animals in the $TUmRNA_6$ group died between days 24 and 25 post-infection, and two animals in the PBS group died between days 25 and 26 post-infection. Therefore, values at subsequent time points in both groups are based on the remaining animals (n = 6). **B)** Guinea pig lung histopathology following $TUmRNA_6$ or $TUmRNA_9$ vaccination and *M. tuberculosis* challenge. The figure shows one representative H&E-stained lung section per group.
