## Supplementary Tables for "Multivalent mRNA tuberculosis vaccines protect against mycobacterial infection in zebrafish, mice and guinea pigs"

**Table S1. The 24 *M. marinum* genes tested for their protective effect in zebrafish. Selection criteria:** Genes were selected for screening based on their differential expression in hypoxic and aerated *M. marinum* cultures, as described in Niskanen et al., 2020[1]. **Gene fragment cloned:** Full-length or partial gene sequences were cloned into DNA vaccines. **Cloning primers:** primers used to amplify selected gene sequences from the genomic DNA of the *M. marinum* ATCC 927 strain.

| M. marinum gene | selection criteria | gene fragment cloned (nucleotides) | Cloning primer (Forward) | Cloning primer (Reverse) |
|---|---|---|---|---|
| MMAR_0010 | upregulated in 2h VS log growth | 166-414 | CTATCTGCTAGCCACCATGCGCTTGGCCCGCGA | CACACAGAATTCGGACGTCAACGCTGGGTGG |
| MMAR_0296 | upregulated in 2h VS 0h | 592-912 | CTATCTGCTAGCCACCATGCGCAAGCCCAAGGTGAGTT | CACACAGAATTCtGCTGGATATCACCACCCGCA |
| MMAR_0678 | upregulated in 2h VS log growth | 1-321 | TCTACAGCTAGCCACCATGATGGGCCTGCACGAGTCG | CACACAGAATTCGTAGTCCGATTCCGCCCAA |
| MMAR_0679 | upregulated IN 2h VS log growth | 160-369 | TCTACAGCTAGCCACCATGATGCTCGCTGTGATCGGCGAT | CACACAGAATTCCCGTCGCGGTTTCGTCGC |
| MMAR_0912 | upregulated in 2h VS 0h+12h VS 0h | 502-849 | CACACTGCTAGCCACCATGGCCAACAGGCTGAAGAC | CACACTGAATTCCGGCGATACAGGAAATCAG |
| MMAR_0913 | upregulated in 2h VS log growth | 1-144 | TCTACAGCTAGCCACCATGCCACACATCGAACAG | TCTTCAGAATTCAGTCGTGCCTGATGACTGATC |
| MMAR_0957 | upregulated in 0h VS M | 43-660 | CACACAGCTAGCCACCATGTCCACCACGGTGCTCGAAG | CACACAGAATTCAATGGGCGAAACCGACAC |
| MMAR_1905 | upregulated in 2h VS 0h | 534-840 | CTATCTGCTAGCCACCCTGCTGCCTGGACCGCTGTGGCAAG | CACACAGAATTCTGTCAACGACAGAGGATCCCGA |
| MMAR_1907 | upregulated in 2h VS 0h+12h VS 0h | 214-516 | CTATCTGCTAGCCACCATGGACGACCACCTGCTGATCC | CACACAGAATTCTGACCTGCGATAGCAGCGC |
| MMAR_2908 | upregulated in 0h VS M | 142-552 | CACACAGCTAGCCACCATGCAAGTGGAGGTTGTCTTC | CACACAGAATTCAGATGGTTCTGTCGTTGTAGAG |
| MMAR_3089 | upregulated in 2h VS 0h | 684-1185 | CTATCTGCTAGCCACCATGAGTCAAACCCATGCTGCAG | CACACAGAATTCACAAGGTGATGTGCAGGTTC |
| MMAR_3206 | upregulated in 2h VS 0h | 10-273 | TCTACAGCTAGCCACCATGATGAGCTCTCCCCCACTTTCAG | TCTTCAGAATTCTCTGGTAGTTGGAGACCGC |
| MMAR_3345 | upregulated in 0h VS M+2h VS log and 12h VS log | 217-963 | CACACAGCTAGCCACCATGGAGAAGGGCACCGATGC | CACACAGGTACCAAGAAGGTCGCCAGCGTGAG |
| MMAR_3642 | upregulated in 2h VS 0h | 1216-1474 | CTATCTGCTAGCCACCATGGTGTTGGGATCGGTATTGG | CACACAGAATTCTCAGAAACACGAAGCCTAGTG |
| MMAR_3673 | upregulated in 2h VS 0h | 1-405 | CTATCTGCTAGCCACCATGGTTGGCGCTTCTGGG | CACACAGAATTCTGGTCTTGCCGATGATCACC |
| MMAR_3806 | upregulated in 2h VS 0h | 1261-1569 | TCTACAGCTAGCCACCATGAGATGACCTGCCCGATC | TCTTCAGAATTCAGGCGAGTTGGCTCTCGT |
| MMAR_3839 | upregulated in 2h VS 0h | 118-270 | TCTACAGCTAGCCACCATGCCAAGGGCTACTGCCCTG | TCTTCAGAATTCGTGGTTCGTCGCCACC |
| MMAR_3841 | upregulated in 2h VS 0h | 1-303 | TCTACAGCTAGCCACCATGGCCAAACATCTAGCG | TCTTCAGAATTCGTGGTTCGTCGCCACC |
| MMAR_4307 | upregulated in 2h VS 0h | 64-384 | CTATCTGCTAGCCACCATGACCAAGCGGCTGCTGGAC | CACACAGAATTCTGGTTTTGGCGATTCGCTC |
| MMAR_4525 | upregulated in 2h VS 0h | 37-376 | CTATCTGCTAGCCACCATGAATATCGTCGACCTGGCCG | GTGTGTGAATTCTGTTCGTCTGTGCGAGCAAGTC |
| MMAR_5144 | Genes expressed exclusively during reactivation, ave >0,125 RPKM | 1-120 | CACACAGCTAGCCACCATGATGAACAACATTTTTCGGGT | CACACAGAATTCTGATGGCGCCGAACGCGGCCG |
| MMAR_5177 | Genes expressed exclusively during reactivation, ave >0,125 RPKM | 16-165 | CACACAGCTAGCCACCATGACCGGGAATAGCGTCGTG | CACACAGAATTCTGATCCCCACCTGCCCGAT |
| MMAR_5324 | Genes expressed exclusively during reactivation, ave >0,125 RPKM | 25-189 | CACACAGCTAGCCACCATGACGCCGCAGGAAGTATTC | CACACAGAATTCTCAGGTCGTCGAGCAACTT |
| MMAR_5410 | upregulated in 2h VS 0h | 208-537 | CTATCTGCTAGCCACCATGCAGGCGCACCAGATTGTG | GTGTGTGAATTCTGGTGGCCAACTCCGAGGT |

[1]Niskanen, M., Myllymäki, H., and Rämet, M. (2020). DNA vaccination with the Mycobacterium marinum MMAR_4110 antigen inhibits reactivation of a latent mycobacterial infection in the adult Zebrafish. Vaccine 38, 5685–5694.
https://doi.org/10.1016/j.vaccine.2020.06.053.

**Table S2. Primers used to amplify *M. tuberculosis* gene sequences from genomic DNA of H37Rv strain**

| ***M. tuberculosis* gene** | **Cloning primer (Forward)** | **Cloning primer (Reverse)** |
|---|---|---|
| Rv0359 | CACACAAGATCTCCACCAtgagcgagacgggccag | CACACATCTAGTtcagaaccagatactccagaa |
| Rv1886c | CACACAAGATCTCCACCatgacagacgtgagccgaa | CACACATCTAGTtcagccggcgcctaacg |
| Rv1234 | CACACAAGATCTCCACCatgactagcccattccagccc | CACACATCTAGTtcagatcgccagacgcg |
| Rv1195 | CACACAAGATCTCCACCAtgtctttcgtgatggcataccca | CACACAACTAGTttagctggccgccgccg |
| Rv2450c | CACACAAGATCTCCACCAtgaagaacgcccgtacgacgct | CACACAACTAGTtcagccgcggcggccgc |
| Rv1386 | CACACAAGATCTCCACCAtgacgttgcgagtcgttc | CACACATCTAGTtcatagcccaccgctga |
| Rv1009 | CACACATGTACAATGttgcgcctggtagtcggtgc | CACACAACTAGTtcagcgcgcacccgctcgtg |
| Rv3875 | CACACAAGATCTCCACCatgacagagcagcagtggaatttc | CACACAACTAGTctatgcgaacatcccagtg |
| Rv2289 | CACACAAGATCTCCACCAtgcccaaaagccgacgg | CACACAACTAGTtcaggacttggtgatggagc |

**Table S3. Reagents used for flow cytometry cell staining.** Related to Figures 3, S4, S5.

| **Target/reagent** | **Fluorochrome conjugate** | **Antibody clone** | **Manufacturer** |
|---|---|---|---|
| BD Horizon™ Fixable Viability Stain 780 | FVS780 | n/a | Waters Biosciences |
| B220 | Alexa Fluor 700 | RA3-6b2 | Waters Biosciences |
| CD44 | APC | IM7 | Waters Biosciences |
| CD3 | PE-Cy7 | 17A2 | Waters Biosciences |
| IgM | PE-CF594 | R6-60.2 | Waters Biosciences |
| CD25 | PE | PC61 | Waters Biosciences |
| CD4 | FITC | RM4-5 | Waters Biosciences |
| CD62L | BV786 | MEL-14 | Waters Biosciences |
| CD8 | BV605 | 53-6.7 | Waters Biosciences |
| PD-1 | BV421 | RMP1-30 | Waters Biosciences |
| CD69 | BV510 | H1.2F3 | Waters Biosciences |
| Mouse BD Fc Block™ | n/a | 2.4G2 | Waters Biosciences |

**Table S4. Antibodies used for flow cytometry cell staining.** Related to Figures S9, S10, S11, S14.

| Target marker | Fluorophore conjugate | Antibody clone | Manufacturer |
|---|---|---|---|
| CD45 | APC-Cy7 | 30-F11 | Waters Biosciences |
| CD127 | PE-Cy7 | SB/199 | Waters Biosciences |
| CD279 | PE | J43 | Waters Biosciences |
| CD3e | AF700 | 500A2 | Waters Biosciences |
| CD3e | UV795 | 145-2C11 | Waters Biosciences |
| CD4 | BV510 | RM4-5 | Waters Biosciences |
| CD44 | FITC | IM7 | Waters Biosciences |
| CD62L | BV786 | MEL-14 | Waters Biosciences |
| CD69 | BV605 | H1.2F3 | Waters Biosciences |
| CD8a | PE-Cy7 | 53-6.7 | Biolegend |
| CD8a | PerCP-Cy5.5 | 53-6.7 | Waters Biosciences |
| KLRG1 | APC | 2F1 | Waters Biosciences |
| NK1.1 | BV421 | PK136 | Waters Biosciences |
| IFN-y | PE | XMG1.2 | Waters Biosciences |
| IL-17A | UV395 | TC11-18H10 | Waters Biosciences |
| IL-2 | BV421 | JES6-5H4 | Waters Biosciences |
| TNF | APC | MP6-XT22 | Waters Biosciences |
